# Chimera-suppressing error-corrected sequencing enables quantitative detection of rare somatic structural-variant breakpoints in human cells

**DOI:** 10.1101/2024.08.08.607188

**Authors:** Elizabeth Pan, Shixiang Sun, Johanna Heid, Zhenqiu Huang, Moonsook Lee, Gordon Fan-Huang, Xiao Dong, Cristina Montagna, Jan Vijg, Alexander Y. Maslov

## Abstract

Rare genome structural-variants (SVs) in somatic cells and tissues are difficult to measure in bulk DNA because library preparation can create chimeric junctions that mimic authentic events. Based on our previous Single-Molecule Mutation sequencing (SMM-seq) assay, we developed a new method, SMM-SV-seq, which combines a chimera-suppressing library preparation with duplex confirmation of breakpoint-supporting molecules. Using libraries from mixtures of human and Drosophila DNA, we show a remaining artifact rate of 1.52 ± 0.53 detectable cross-species chimeras per billion bases, providing an empirical background for correcting breakpoint frequencies. In a benchmark of evaluable homozygous germline deletions in the GM24385 cell line, SMM-SV-seq achieved 81.4% precision and 85.7% recall, exceeding the precision of established short-read callers while maintaining comparable recall. The assay detected dose-associated increases in SV-breakpoint frequencies in cells treated with bleomycin, a known clastogen. Biological relevance was demonstrated by a significantly greater irradiation-associated increase in translocation-like breakpoint frequencies in the BRCA1 185delAG-heterozygous mammary epithelial clone than in its isogenic wild-type control. These results establish SMM-SV-seq as a practical assay for calibrated, duplex-confirmed detection of rare somatic SV breakpoints directly in bulk DNA from non-clonal cell populations.

---

Somatic mutations contribute to cancer and are increasingly implicated in aging and other diseases^1–3^. Rare somatic mutations are difficult to detect by conventional bulk sequencing because most de novo mutations in normal cells or tissues are confined to individual cells or small clones. Advances in single-molecule, error-corrected next-generation sequencing (ecNGS) now enable measurement of somatic mutagenesis in genetic toxicology testing and research in cancer and aging^4,5^. However, most ecNGS workflows have been developed for detecting single-nucleotide variants (SNVs) and short insertions or deletions (INDELs). Their application to rare somatic SV-associated breakpoints is limited by double-stranded chimeric junctions created during library preparation, which can survive duplex consensus filtering and mimic authentic breakpoints^6^. Of note, PCR amplification is another source of chimeric artifacts^7^, but these arise after strands are labeled and are efficiently removed by duplex error correction because they are exceedingly unlikely to recur identically on both strands of the same molecule.

The difficulty of reliably detecting somatic SVs is a major limitation in studying genome instability in relation to cancer and aging because SVs can have greater functional consequences than SNVs or small INDELs^8^. SVs represent a highly heterogeneous class of mutations, arbitrarily defined as events involving more than 50 bp, including everything from deletions to chromosomal translocations. Their accurate detection is further complicated by their substantially lower frequency relative to single-nucleotide variants. Population-scale short-read sequencing studies initially identified approximately 2,000–2,500 germline SVs per human genome^9^. Improved detection using long-read sequencing has since revealed approximately 25,000 SVs per diploid genome, still far fewer than the approximately four million single-nucleotide variants^10^. Quantitative data on the frequency and spectrum of somatic SVs in normal cells remain limited^8^.

Here we describe SMM-SV-seq, a short-read assay that quantifies rare somatic SV-associated breakpoint frequencies by combining Tn5-mediated, chimera-suppressing library preparation with rolling-circle amplification and duplex error correction. Breakpoint detection was benchmarked using a curated set of previously annotated homozygous deletions in the GM24385 cell line^11^, restricted to autosomal loci covered by duplex fragments. Libraries prepared from mixtures of human and Drosophila DNA were used to estimate a total residual-chimera rate of approximately 3 junctions per billion qualifying duplex bases. We show that the method detects dose-associated increases in SV-breakpoint frequencies after exposure to bleomycin, a known clastogen, and a greater irradiation-associated increase in translocation-like breakpoint frequencies in the tested BRCA1 185delAG-heterozygous clone than in its isogenic wild-type control.

## Results

### Chimera-suppressing SMM-SV-seq workflow

To enable reliable detection of somatic SV-associated breakpoint junctions, we redesigned the SMM-seq library preparation to suppress pre-labeling chimeric junction formation (Fig. 1A). DNA fragmentation and library construction are now performed using Tn5-mediated transposition, followed by a dedicated end-processing step that generates defined overhangs complementary to those on the hairpin adapters and enforces strict ligation compatibility (Fig. 1B). This design is intended to suppress illegitimate fragment–fragment ligation while preserving the dumbbell structures required for rolling circle amplification (RCA). The resulting circular templates are amplified by RCA and processed using duplex error correction, retaining the ability to filter post-labeling errors that lack concordant support from both complementary strands. Together, ligation suppression during library construction and duplex consensus filtering in the analysis enable confident detection of SV-associated breakpoints from individual duplex molecules.

**Figure 1.**
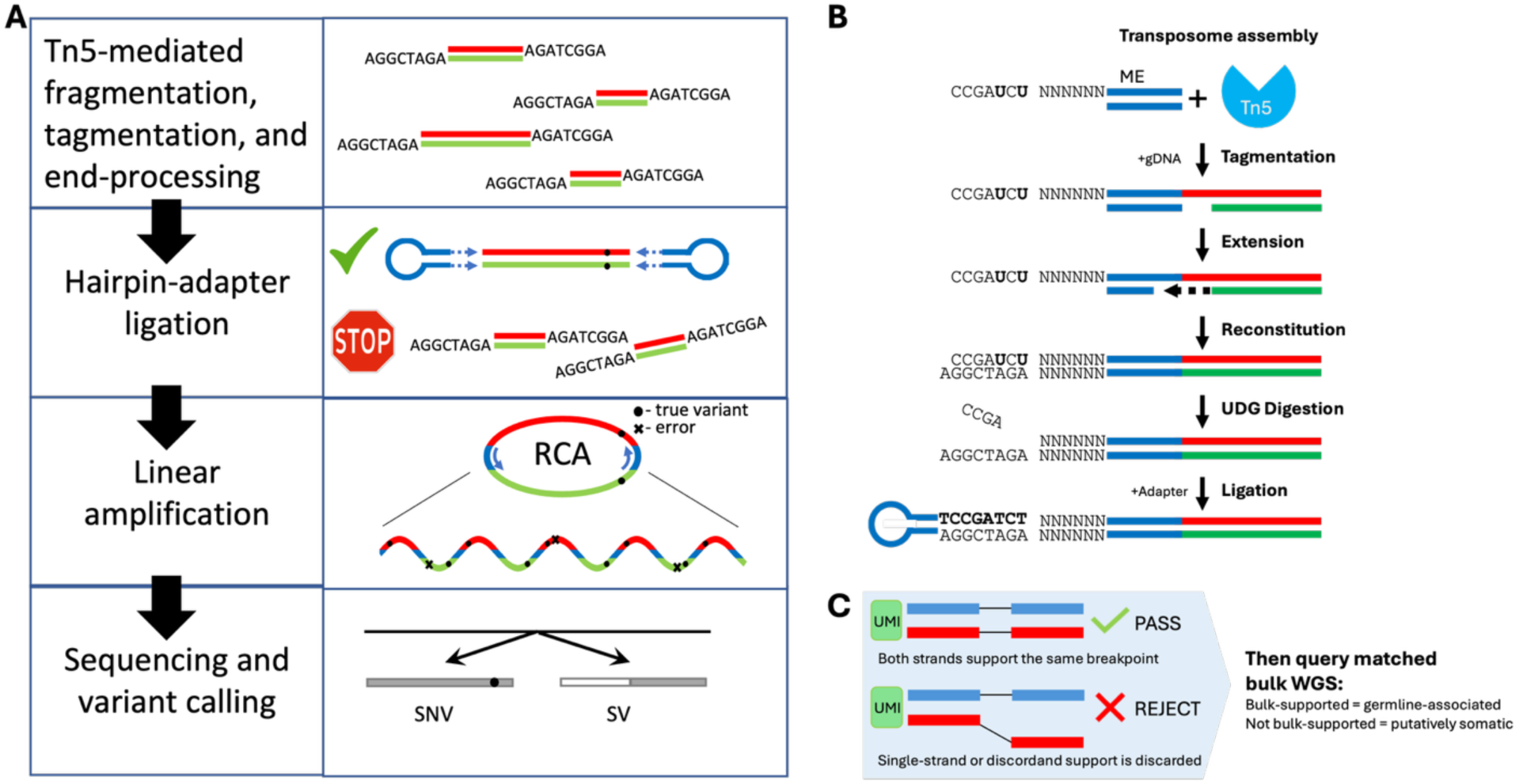
Chimera-suppressing SMM-SV-seq library preparation and duplex-supported breakpoint-junction classification. (**A**) Overview of the SMM-SV-seq workflow. Genomic DNA is fragmented and tagged using hyperactive Tn5 transposase. Following end processing, hairpin adapters are ligated to compatible cohesive ends, while fragment–fragment ligation is disfavored. Hairpin-ligated molecules undergo rolling-circle amplification (RCA), and products derived from the complementary strands of each source molecule are sequenced and analyzed independently. Filled circles indicate variants present in the source molecule, whereas crosses indicate errors introduced after strand labeling. (**B**) Detailed view of transposome assembly and end processing. The upper transposon oligonucleotide contains two deoxyuridines, a six-nucleotide random unique molecular identifier (UMI), and the Tn5 mosaic-end (ME) sequence. Following tagmentation, DNA-polymerase extension repairs the staggered products and copies the UMI to the complementary strand. USER treatment cleaves at the deoxyuridines, exposing defined 8-nucleotide 3′ overhangs. Hairpin adapters carrying complementary overhangs are then ligated with T7 DNA ligase, favoring compatible cohesive-end ligation over blunt-end fragment–fragment ligation. (**C**) Simplified duplex-support and bulk-annotation logic. A candidate SV-associated breakpoint junction is retained when the two complementary source strands support the same reciprocal discordant mapping configuration; single-strand support or disagreement between the complementary strands is rejected. Retained junctions are queried against matched conventional whole-genome sequencing (WGS): junctions with bulk support are classified as germline-associated, whereas junctions without bulk support are classified as putatively somatic. Red and green lines represent the complementary genomic DNA strands, and blue elements represent transposon and hairpin-adapter sequences. N, random nucleotide; gDNA, genomic DNA. Schematics are not to scale.

### Duplex-informed SV breakpoint calling and germline annotation

SMM-SV-seq data analysis leverages both strand identity and mapping concordance to identify SV-associated breakpoint junctions (Fig. 1C; Supplementary Fig. S1A). Sequencing read pairs are grouped by unique molecular identifiers (UMIs) and alignment-start coordinates, and complementary source strands are identified after deduplication. Candidate SV-associated breakpoint junctions are retained only when the two complementary strands support reciprocal mirrored discordant mapping configurations. Based on genomic alignment patterns, discordant read pairs mapping to the same chromosome with anomalous spacing are annotated as deletion-like junctions, whereas pairs mapping to different chromosomes are annotated as translocation-like junctions^12,13^.

To classify candidate breakpoint junctions as germline-associated or putatively somatic, each duplex-confirmed breakpoint is queried in conventional bulk sequencing data generated from the same sample (Supplementary Fig. S1A). Although conventional libraries contain abundant chimeric background, the likelihood that an independent artifactual junction coincides with the exact breakpoint detected by SMM-SV-seq is exceedingly low. Accordingly, candidate junctions supported by independently generated read pairs in bulk sequencing data are classified as germline-associated, whereas those lacking such support are designated putatively somatic. Putative somatic breakpoint frequencies are quantified relative to the total genomic region covered by proper duplex fragments (Supplementary Fig. S1B).

### Benchmarking SMM-SV-seq against established SV callers for bulk DNA

We next benchmarked the accuracy of SMM-SV-seq against a curated set of known deletions in the well-characterized GM24385 cell line (Fig. 2). The GM24385 autosomal reference set contained 771 deletions ≥1 kb, of which 234 are reported as homozygous and 537 as heterozygous. Because SMM-SV-seq interrogates DNA molecules by random sampling, only breakpoints located within genomic regions covered by duplex fragments were considered evaluable. Importantly, covered homozygous deletions have a higher expected probability of detection than heterozygous deletions and therefore provide an appropriate benchmark set. This setup provided a robust dataset for comparing SMM-SV-seq with established computational SV detection tools, including Manta^14^, DELLY^15^, and GRIDSS^16^.

**Figure 2.**
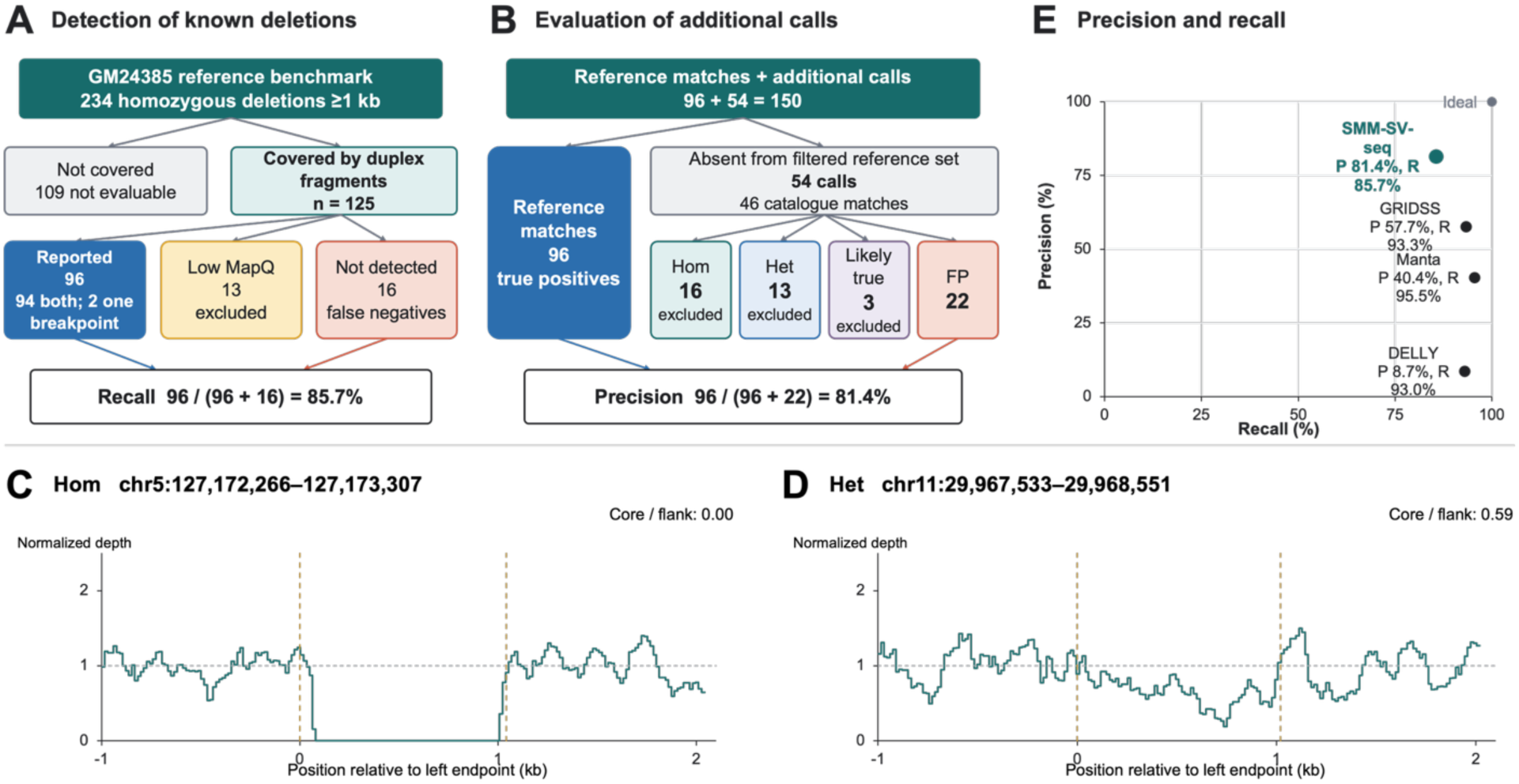
Germline deletion benchmarking and representative coverage profiles. (**A**) Classification of autosomal reference homozygous deletions ≥1 kb by duplex-fragment coverage and SMM-SV-seq detection. Uncovered loci and calls removed by mapping-quality filtering were excluded from recall. (**B**) Classification of reference matches and additional calls. For precision, 16 homozygous (Hom), 13 heterozygous (Het) and three likely true deletions were excluded from both the true-positive and false-positive counts; 22 calls were conservatively counted as false positive (FP). “Likely true” denotes deletions with supporting catalogue and bulk WGS evidence but unresolved genotype. The 96 reference matches correspond to 94 unique SMM-SV-seq intervals; the total of 150 represents reference matches plus 54 additional calls. (**C,D**) Representative PASS homozygous (C, genotype 1/1) and heterozygous (D, genotype 0/1) deletions from the full GM24385 catalogue. Coordinates are GRCh37/hg19. Teal curves show coverage from the bulk WGS alignment. Only primary alignments with MAPQ ≥20 were counted; duplicates and QC-fail alignments were excluded. Mean depth was calculated in 200 bins across the reported interval and its two 1-kb flanks, then normalized to the arithmetic mean of the unbinned left– and right-flank depths. Dashed vertical lines mark the reported interval; the dashed horizontal line marks normalized depth of 1. Core/flank ratios use mean unbinned interval depth after trimming 200 bp from each end and the same flank normalization. (**E**) Precision and recall at the reported operating point for each caller. Ideal denotes 100% precision and 100% recall. No inferential statistical comparison was performed.

Among the 234 homozygous deletions annotated in GM24385, 125 were covered in the SMM-SV-seq data; the remaining 109 were not covered and were excluded from evaluation. Of the 125 covered deletions, 96 were reported by SMM-SV-seq (Fig. 2A; Supplementary Table 1). Of the covered set, 13 deletions were detected by SMM-SV-seq but failed a conservative mapping-quality filter and were therefore not reported. These 13 deletions were excluded from the recall calculation. The remaining 16 covered deletions had no matching call and were counted as false negatives. Under this exclusion rule, recall was 85.7% [96/(96 + 16)].

Each deletion is bounded by two breakpoints. Of the 96 reported deletions, 94 matched the reference at both breakpoints and two matched at only one breakpoint. For 86 of the 94 two-breakpoint matches (91.5%), both breakpoint positions were within 250 bp of the reference coordinates (Supplementary Fig. S2).

SMM-SV-seq also reported 54 recurrent deletions not present in the size-filtered reference set of deletions ≥1 kb (Fig. 2B; Supplementary Table 2). However, comparison with the full GM24385 deletion catalogue identified matches for 46 of these 54 calls. Twenty-two matched PASS entries: nine homozygous deletions (Fig. 2C) and 13 heterozygous deletions (Fig. 2D). All 22 PASS entries had annotated lengths shorter than 1 kb and had therefore been excluded from the size-filtered reference set. Among the remaining 32 calls, coverage loss in conventional whole-genome sequencing data supported seven additional homozygous deletions. Six of these seven calls matched non-PASS catalogue entries. Three further calls also matched non-PASS catalogue entries at both breakpoints, but showed partial coverage loss with multiple high-quality supporting bulk WGS read pairs. These were considered likely true deletions with unresolved genotype. Under the reference-based scoring scheme, 32 of the 54 calls were excluded from both the true-positive and false-positive counts used to calculate precision: 16 homozygous deletions (nine PASS catalogue matches and seven additional coverage-supported calls), 13 heterozygous deletions and three likely true deletions with unresolved genotype. The remaining 22 were conservatively counted as false positives, although supporting read pairs were observed in the bulk WGS dataset. Of note, 15 of these 22 calls also matched non-PASS catalogue entries. Under this scoring scheme, precision was 81.4% (96 / (96 + 22)).

In this benchmark, SMM-SV-seq precision (81.4%) exceeded that of Manta (40.4%), DELLY (8.7%), and GRIDSS (57.7%). Recall was 85.7% for SMM-SV-seq, compared with 95.5% for Manta, 93.0% for DELLY, and 93.3% for GRIDSS (Fig. 2E).

### Evaluation of false-positive rate

To directly quantify the rate of possible residual library-derived chimeric artifacts, we performed a species-mixing control experiment. Human genomic DNA was mixed at a 1:1 molar ratio with *Drosophila melanogaster* genomic DNA and processed through the complete SMM-SV-seq workflow, with reads aligned to a combined human-fly reference genome. In this experimental design, any read pair in which one mate maps to the human genome and the other to the fly genome must result from artifactual ligation during library preparation, as no true structural variants can connect sequences from the two species (Fig. 3A).

**Figure 3.**
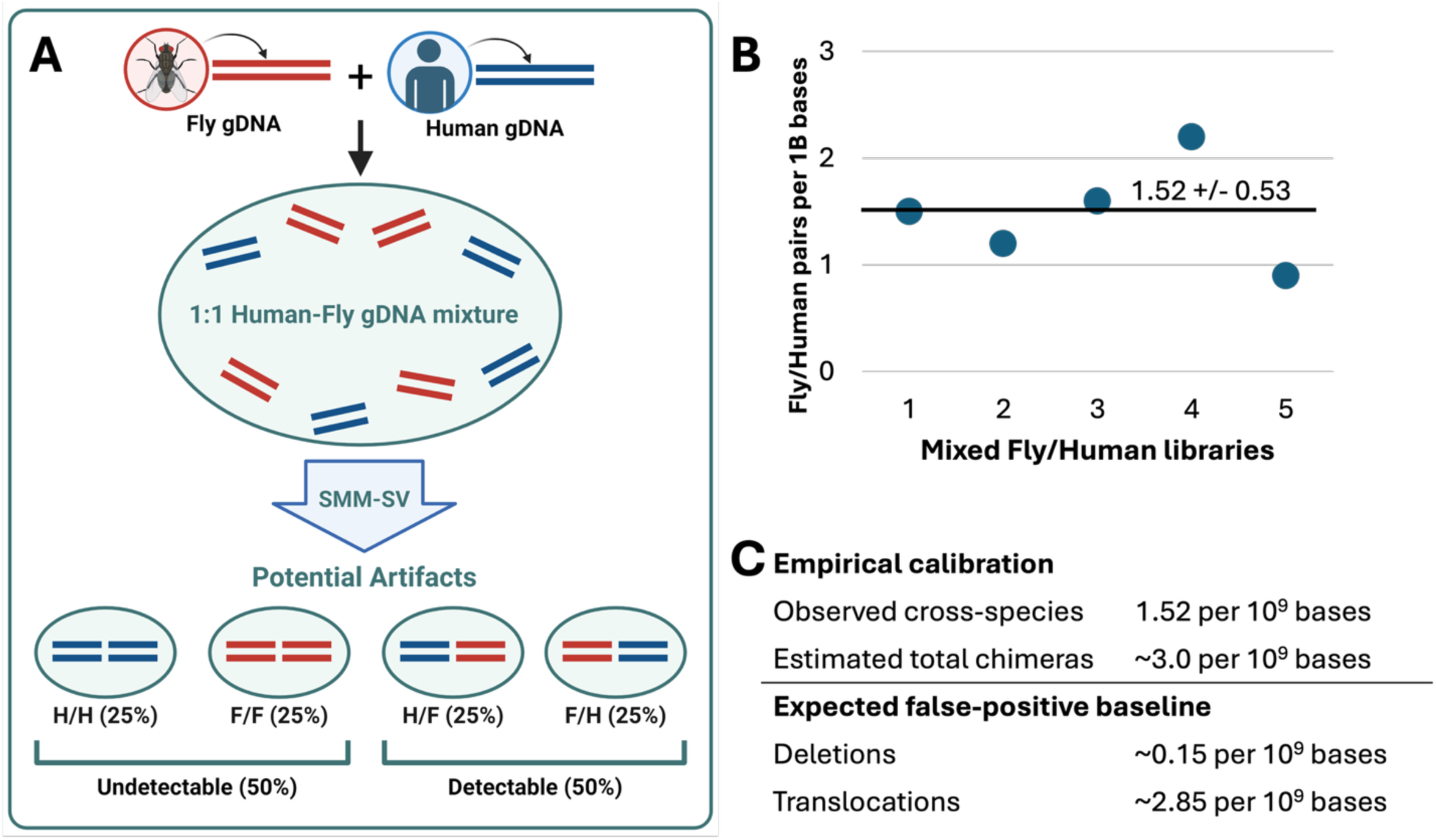
Species-mixing calibration of residual library-derived chimera background. (**A**) Human and Drosophila melanogaster genomic DNA were mixed at a 1:1 molar ratio and processed using SMM-SV-seq. Assuming random ligation, cross-species human–fly and fly–human junctions represent 50% of library-derived chimeras and are directly detectable; same-species junctions cannot be distinguished from intragenomic events. (**B**) Cross-species junction rates in five independently prepared mixed-DNA libraries. Each point represents one library. Rates were normalized to the cumulative length of qualifying proper-duplex fragments and are reported per 10⁹ bases. The mean observed rate was 1.52 ± 0.53 cross-species junctions per 10⁹ bases. (**C**) Doubling the observed cross-species rate produced an estimated total residual-chimera rate of approximately 3.0 per 10⁹ bases. Assuming random ligation and an expected distribution of approximately 5% intrachromosomal and 95% interchromosomal junctions, the estimated false-positive baselines were 0.15 deletion-like and 2.85 translocation-like junctions per 10⁹ bases, respectively. These values were subtracted from the corresponding rates in subsequent analyses.

Sequencing and analysis of five independently prepared mixed-species libraries in two separate experiments identified approximately nine duplex-supported cross-species junctions per library, with each library containing an average of 16.5 million duplex consensus molecules (i.e., double-stranded DNA fragments). In contrast, when libraries prepared from human-only or fly-only DNA were aligned to the same combined human-fly reference, we observed no read pairs with mates mapping to different species, supporting mapping specificity under these control conditions.

Across the five mixed-species libraries, the frequency of detectable cross-species junctions was 1.52 ± 0.53 per 10⁹ qualifying duplex bases (Fig. 3B), providing a direct empirical estimate of the observable residual chimera rate in the SMM-SV-seq workflow. Importantly, in a 1:1 species mixture, only half of all ligation-derived chimeric artifacts are expected to be detectable as cross-species events; the remaining half arise from human–human or fly–fly ligation products and are therefore indistinguishable from genuine intragenomic structural variants. Accordingly, the estimated false-positive baselines were approximately 0.15 deletion-like and 2.85 translocation-like junctions per 10⁹ qualifying duplex bases (Fig. 3C).

To extrapolate these findings to a pure human DNA context, we considered the chromosomal organization of the human genome. Assuming random ligation between genomic fragments, approximately 5% of artifactual junctions are expected to occur within the same chromosome (intrachromosomal), while the remaining ∼95% will involve fragments from different chromosomes (interchromosomal). Accordingly, using the total artifact rate of ∼3 per billion bases, the estimated false-positive rates are ∼0.15 per billion bases for deletions and ∼2.85 per billion bases for translocations.

Together, these analyses provide an empirical estimate of the residual chimera background under the tested conditions. This estimate was used to correct downstream deletion-like and translocation-like breakpoint frequencies.

### Dose-associated increases in somatic SV-breakpoint frequencies after clastogenic stress in primary human cells

Having established a low false-positive baseline and benchmarked performance, we next evaluated the utility of SMM-SV-seq for genetic toxicology applications. For this purpose, we used the method to measure somatic SV-associated breakpoint frequencies in human IMR90 primary fibroblasts exposed to bleomycin, a well-characterized radiomimetic clastogen^17^. Cells were exposed to 0, 5, 10, or 20 μM bleomycin and collected 72 h later, with three independent biological replicates analyzed per condition; this dose range was compatible with high to modest survival rates^18^.

Across these experiments, SMM-SV-seq achieved an average genomic coverage of approximately 3 billion bases per sample. Matched conventional sequencing libraries generated from the same cell populations were used to identify and filter germline SVs, enabling quantitative analysis of somatic SV-associated breakpoint frequencies. All reported breakpoint frequencies were corrected for residual library-derived artifacts by subtracting the empirically measured false-positive rate determined in the species-mixing experiment described above.

Bleomycin exposure was associated with increases in corrected deletion-like and translocation-like breakpoint-junction frequencies relative to untreated controls (Fig. 4A,B). In IMR90 primary human fibroblasts collected 72 h after treatment, mean deletion-like frequencies were 1.4 ± 0.2, 4.2 ± 0.7, 5.3 ± 1.4, and 10.8 ± 2.6 per 10^9^ bases covered by proper duplex fragments at 0, 5, 10, and 20 μM bleomycin, respectively. The corresponding translocation-like frequencies were 14.1 ± 1.4, 56.2 ± 27.4, 95.2 ± 10.8, and 155.7 ± 27.6 per 10^9^ covered bases. Thus, SMM-SV-seq detected dose-associated increases in SV-associated breakpoint-junction frequencies in primary human fibroblasts under the tested conditions.

**Figure 4.**
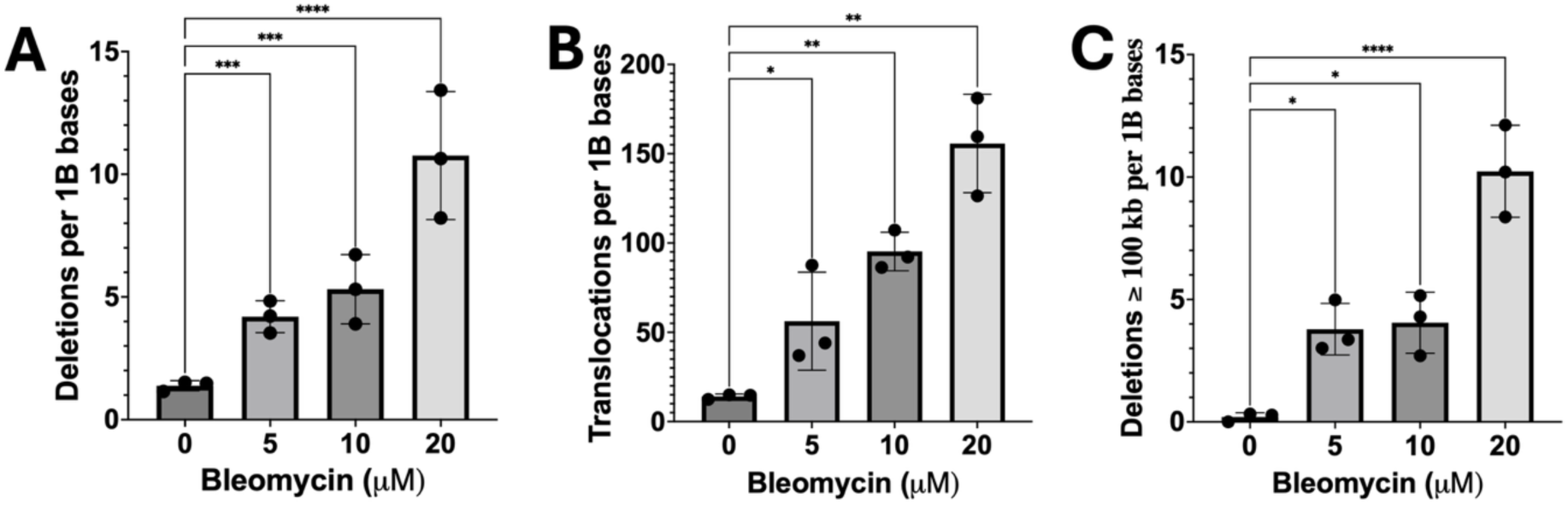
Bleomycin-associated changes in SV-associated breakpoint-junction frequencies in IMR90 fibroblasts. Corrected deletion-like (**A**) and translocation-like (**B**) junction frequencies and the frequency of deletion-like junctions with paired breakpoints separated by at least 100 kb in the reference genome (**C**), measured 72 h after exposure to the indicated bleomycin concentrations. Each point represents an independent biological replicate (n = 3 per condition); bars show mean ± s.d. Frequencies are reported per 10⁹ qualifying duplex bases after subtraction of the corresponding background rate. *P < 0.05, **P < 0.01, ***P < 0.001, ****P < 0.0001.

Analysis of deletion-like junction span showed that the bleomycin-associated increase included a prominent long-range component (Fig. 4C). Mean frequencies of deletion-like junctions spanning at least 100 kb were 0.2 ± 0.2, 3.8 ± 1.1, 4.1 ± 1.2, and 10.2 ± 1.9 per 10⁹ qualifying duplex bases at 0, 5, 10, and 20 μM bleomycin, respectively; all three treated groups differed significantly from untreated controls. Across all bleomycin-treated samples, 88.0% (139 of 158) of the deletion-like junctions had inferred spans of at least 100 kb, compared with 13.3% (2 of 15) in untreated controls. Thus, long-span junctions constituted the great majority of deletion-like calls observed after bleomycin exposure. This enrichment is consistent with reporter-locus evidence that bleomycin can induce very large deletions in CHO cells^19^ and cytogenetic evidence of an elevated proportion of interstitial deletions in bleomycin-treated human lymphocytes^20^. Accordingly, we describe these events as long-span deletion-like junctions, with span defined by the reference-coordinate separation between the joined loci.

Together, these results show that SMM-SV-seq detects dose-associated increases in SV-associated breakpoint frequencies in primary human cells. This supports further evaluation of SMM-SV-seq as a breakpoint-level approach for genetic toxicology applications^21^. The assay measures dose-associated changes in breakpoint frequency and may support future studies of breakpoint patterns. Importantly, these features make SMM-SV-seq a potentially valuable assay for evaluating compounds with suspected clastogenic activity.

### Irradiation-associated SV-breakpoint frequencies in BRCA1 185delAG-heterozygous cells

BRCA1 is a major cancer risk gene. Haploinsufficiency of BRCA1 due to germline mutations is associated with an increased risk of cancer, most notably breast and ovarian cancer^22^. While BRCA1 is a key component of homologous recombination (HR) repair, and haploinsufficiency compromises the resolution of DNA double-strand breaks^23^, almost nothing is known about its effect on SV frequency and spectrum. To test whether BRCA1 genotype was associated with a different SV-breakpoint response to irradiation, we applied SMM-SV-seq to human mammary epithelial cells heterozygous for the pathogenic BRCA1 185delAG mutation, derived from primary telomerized mammary epithelial cells (hTERT-IMECs)^24^. An isogenic wild-type (WT) hTERT-IMEC line served as the matched control. Both cell lines were analyzed without irradiation and after a single 2 Gy dose of IR, with three independent biological replicates per genotype and treatment condition. SV-associated breakpoint frequencies were normalized per 10^9^ bases covered by proper duplex fragments and corrected for residual library-derived artifacts.

The BRCA1 185delAG-heterozygous line showed a larger post-irradiation increase in reported SV-associated breakpoint frequencies than the isogenic WT line (Fig. 5). In WT cells, deletion-like breakpoint frequencies increased from 1.4 ± 0.6 in untreated cells to 8.0 ± 2.7 per billion bases after irradiation, whereas in the BRCA1 185delAG-heterozygous line deletions increased from 6.4 ± 3.8 to 18.0 ± 8.1 per billion bases (Fig. 5A). A similar genotype-dependent pattern was observed for translocation-like breakpoint frequencies which increased from 16.9 ± 8.8 to 280.1 ± 199.0 per billion bases in WT cells and from 72.8 ± 3.8 to 633.6 ± 96.6 per billion bases in the BRCA1 185delAG-heterozygous line (Fig. 5B).

**Figure 5.**
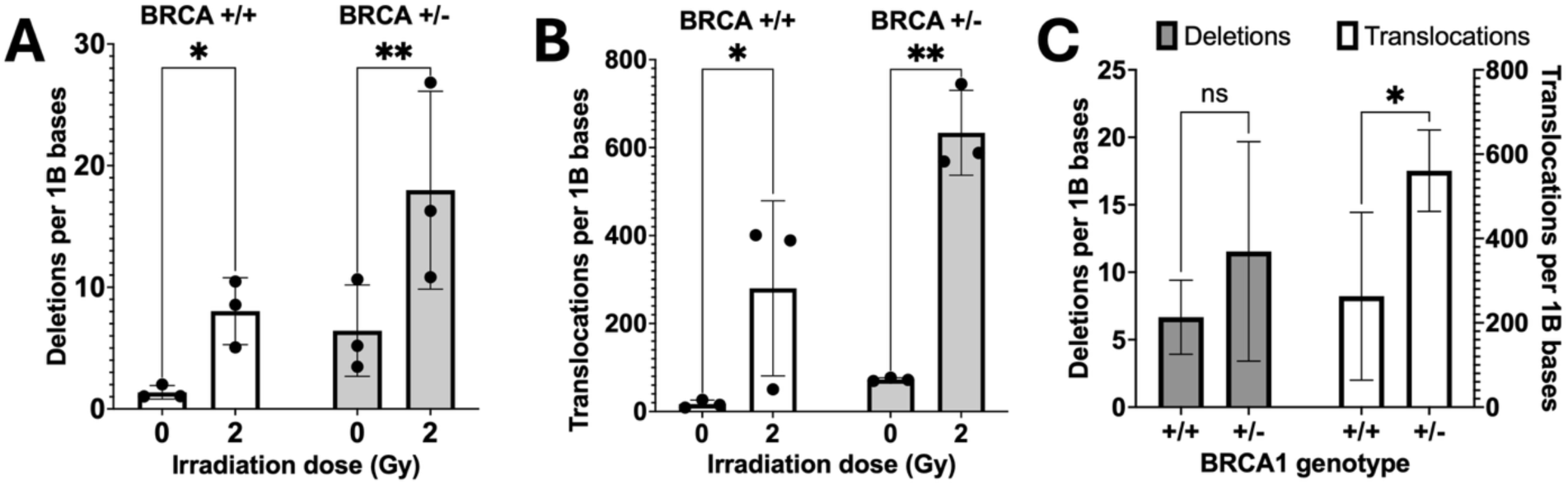
Irradiation-associated SV-breakpoint frequencies in the tested BRCA1 185delAG-heterozygous clone and its isogenic wild-type line. Corrected deletion-like (**A**) and translocation-like (**B**) junction frequencies in BRCA1+/+ and BRCA1 185delAG/+ hTERT-IMEC cells before and 72 h after 2 Gy irradiation. (**C**) Irradiation-associated changes from baseline in deletion-like and translocation-like junction frequencies compared between the BRCA1 185delAG/+ clone and its isogenic wild-type line. Each point in A and B represents a biological replicate from three matched experiments; bars show mean ± s.d. Frequencies are reported per 10⁹ qualifying duplex bases after subtraction of the corresponding background rate. *P < 0.05, **P < 0.01.

Although baseline breakpoint frequencies differed between the lines before irradiation, potentially reflecting clonal history, the absolute post-irradiation increases were numerically larger in the BRCA1 185delAG-heterozygous line: 11.5 versus 6.7 deletion-like junctions and 560.8 versus 263.2 translocation-like junctions per 10^9^ bases covered by proper duplex fragments (Fig. 5C). The between-line difference was significant for the translocation-like response but not for the deletion-like response. This result identifies a difference in the translocation-like response between the tested BRCA1 185delAG-heterozygous clone and its isogenic wild-type control but does not identify the responsible mechanism, because homologous-recombination activity and repair-pathway use were not measured directly. These data demonstrate that SMM-SV-seq can compare irradiation-associated breakpoint-frequency responses across defined cultured human-cell models.

## Discussion

Quantifying somatic mutations in normal tissues has long been challenging. Reporter-gene models enabled sensitive detection of somatic SNVs and SVs at engineered loci in vivo^25^, but genome-wide measurement in human cells became feasible only with the emergence of single-cell and single-molecule sequencing approaches^26–28^. However, these approaches have largely been limited to SNVs and small INDELs. Efforts to enable rare somatic SV detection have included SVS, which used single-stranded adapter ligation to suppress chimera formation but lacked complementary-strand confirmation of individual junctions^29^. In targeted svCapture, ligation artifacts persisted despite duplex filtering, whereas its chimera-suppressing tagmentation workflow did not support duplex assignment^6^. In the present study, SMM-SV-seq combines chimera-suppressing library preparation with complementary-strand confirmation to quantify SV-associated breakpoint frequencies without target capture, using empirical calibration of the residual-artifact background. Specifically, junction artifacts are minimized by controlling DNA end structures and enforcing ligation compatibility, with post-labeling errors filtered by consensus-based correction. This layered design targets the main technical failure mode in short-read somatic SV detection, where a small fraction of library-derived junctions can persist through standard processing and inflate apparent SV burden.

The species-mixing control provides an empirical estimate of detectable residual chimera background under the tested library-preparation and analysis conditions. The deletion– and translocation-specific correction values depend on the species-mixture design and the random-ligation assumptions described above. Applying an experimentally measured artifact rate enables quantitative correction of somatic SV-associated breakpoint frequencies, which is important in normal cells where true breakpoint burdens are low and technical noise could otherwise dominate the biological signal. In our experiments, the empirically defined background was below the breakpoint frequencies observed in untreated human cells.

In the absence of a direct gold standard for somatic SVs, germline deletions in a well-characterized human cell line served as a surrogate benchmark because they are detected by breakpoint evidence in the same manner as somatic events. Within the evaluated GM24385 homozygous-deletion benchmark, SMM-SV-seq showed higher precision than Manta, DELLY and GRIDSS under the stated reference-based scoring and exclusion rules, with a recall of 85.7%.

These results support further evaluation of SMM-SV-seq for genetic toxicology and genotype-comparison studies. In bleomycin-treated primary fibroblasts, corrected breakpoint frequencies increased with dose. In the tested BRCA1 185delAG-heterozygous mammary epithelial clone, irradiation was followed by a larger increase in deletion-like and translocation-like breakpoint frequencies than in the isogenic wild-type line. Because repair activity was not measured, these data do not establish impaired double-strand-break resolution or another specific mechanism.

Importantly, SMM-SV-seq reports structural variation at the level of individual junctions rather than attempting to infer complete rearrangement structures from clusters of supporting reads. As emphasized by the SV classification framework of Li et al., many structural variants are inherently complex and can comprise multiple breakpoints organized into nested or compound architectures^30^. SMM-SV-seq is well suited to sensitively detect such constituent junctions, but short-read data generally do not permit reconstruction of the full rearrangement in many cases. This limitation is important for interpreting events classified as deletions and translocations, which in breakpoint-based assays can also represent the footprint of localized insertions, including transposon-mediated events^31^. Integration with long-read platforms (Oxford Nanopore or PacBio) offers a direct route to refine SV classification and reconstruct event architectures^32^.

Overall, SMM-SV-seq addresses a specific measurement gap in bulk DNA: library-derived chimeric junctions can confound quantitative detection of rare somatic SV-associated breakpoints. By combining chimera-suppressing library preparation with single-molecule error correction, SMM-SV-seq extends the SMM-seq workflow to calibrated, quantitative measurement of duplex-supported SV-associated breakpoint-junction frequencies in cultured human cells. Although the residual false-positive rate is low, further improvements in chimera suppression and calibration should reduce background and extend sensitivity to rarer events.

## Materials and Methods

### Cell culture and treatment

Human normal lung IMR90 fibroblasts were cultured under a 10% CO2 and 3% O2 atmosphere at 37°C in DMEM (GIBCO, Grand Island, NY, USA), supplemented with 10% FBS (GIBCO). Human mammary epithelial cells hTERT-IMEC and hTERT-IMEC no. 2 were cultured in MEBM supplemented with Bullet kit (Lonza)^33^. Twenty-four hours post-seeding, the culture media was replaced with media containing various doses of bleomycin and the cells were exposed for three hours. Subsequently, after cells were washed with PBS, the media was replaced with fresh complete media. Cells were harvested 72 hours post-bleomycin exposure. Control cells were maintained in media containing only the vehicle. For irradiation experiments, mammary epithelial cells were irradiated with a single 2 Gy dose of ionizing radiation (IR) 24 hours after plating and harvested 72 hours later.

### DNA isolation and quality control

DNA was isolated from fibroblasts and mammary epithelial cells using the Quick-gDNA™ MiniPrep kit (Zymo Research Corporation, Irvine, CA, USA) following the manufacturer’s instructions. The concentration of DNA was quantified using a Qubit kit (ThermoFisher Scientific, USA). DNA integrity was assessed using the 4200 TapeStation System (Agilent Technologies, Inc., CA, USA), ensuring a DNA Integrity Number (DIN) above 8.0.

### SMM-SV library preparation and sequencing

Unloaded Tn5 hyperactive transposase was sourced from Diagenode. Transposome assembly was conducted according to the manufacturer’s recommendations using the following oligonucleotides (IDT, USA): CCGA/ideoxyU/C/ideoxyU/ NNNNNN AGA TGT GTA TAA GAG ACA G and /5Phos/CTG TCT CTT ATA CAC ATC T. The tagmentation reaction was performed using 100 ng of genomic DNA at 55°C for 7 minutes, following manufacturer guidelines. Tagmentation products were purified using the Clean and Concentrate kit (Zymo), eluted in 24 µL of elution buffer, and mixed with 6 µL of Taq 5X Master Mix (NEB, USA). This mixture was incubated at 65°C for 30 minutes to repair staggered nicks and replace the bottom strand on the introduced transposon tag. Subsequently, DNA fragments were purified with 1.5X AMPure XP beads and resuspended in 23 µL 0.1x TE buffer.

For exposing the 3’-end overhangs, the purified DNA fragments were supplemented with 3 µL T4 ligase buffer, 1 µL of SVFP9_HP adapter (10µM), and 2 µL of USER enzyme (NEB), which digests the uracil bases to expose the 3’-end overhangs. This mixture was incubated at 37°C for 15 minutes. Then, 1 µL of T7 ligase was added, followed by a 5-minute incubation and heat inactivation of T7 ligase at 65°C for 10 minutes. The SVFP9_HP adapter was prepared by heating 20 µL of a 100 µM solution of oligonucleotide /5Phos/ AGA GCA CAC GTC TGA ACT CCA GTC TAC ACT CTT TCC CTA CAC GAC GCT CTT CCG ATC T-3’ (IDT, USA) in 0.1X TE buffer to 95°C for 5 minutes, then cooling to 37°C for 5 minutes to form a hairpin.

The ligated product was treated with exonuclease III (NEB) at 37 °C for 15 min, followed by another round of purification with 1.5× AMPure XP beads and size selection for 600–800 bp fragments using a PippinHT system (Sage Science, USA). Then samples were diluted based on assessed molar concentration as described previously^28^. A 13.4 µL aliquot of the diluted sample was used in a pulse-RCA reaction, performed in a 20 µL volume containing 10U (0.5 µL) of SD polymerase HS (BIORON Diagnostics GmbH, Germany), along with SD polymerase buffer, 10 mM dNTPs mix (NEB), 0.6 µL 100 mM MgCl2, and water. The pulse-RCA program included an initial denaturation at 92°C for 2 minutes, followed by 14 cycles of denaturing, annealing, and extending steps, and a final hold at 4°C. The RCA product was then purified, PCR amplified using NEBNext Ultra II Q5 Master Mix and dual index oligos, and sequenced on an Illumina NovaSeq instrument in 150 paired-end mode. Conventional sequencing libraries were prepared using the NEBNext Ultra II FS DNA Library Prep Kit for Illumina (NEB) according to the manufacturer’s instructions. After quantification with Qubit, samples were pooled and sequenced on Illumina NovaSeq instrument using 150 paired-end mode.

### Read processing, alignment and computational provenance

Human-cell libraries used for the bleomycin and irradiation analyses were processed with archived custom SVD v0.5 scripts. Raw paired-end reads were inspected with FastQC and adapter– and quality-trimmed with Trim Galore v0.4.1. Trimmed reads were aligned to the 1000 Genomes GRCh37 reference with decoy sequences using BWA-MEM v0.7.13-r1126 with the – M option. Alignments were processed using SAMtools v1.9 and Picard v1.119. GATK v3.5-0-g36282e4 was used for local realignment around known indels, using the 1000 Genomes Phase 1 and Mills/1000 Genomes gold-standard indel resources, followed by the archived PrintReads step. Conventional duplicate removal was omitted before UMI-based processing. Conventional libraries used for junction annotation were aligned to the same human reference. Structural-variant analysis was restricted to chromosomes 1–22.

### UMI grouping and complementary-strand identification

The six-nucleotide UMI introduced at each end of a source DNA fragment was treated as an ordered pair of end tags. Read pairs were grouped using the paired UMIs and their alignment-start coordinates, and the longest read pair was retained within each duplicate group. Complementary-strand pairs were identified by reciprocal ordering of the two end UMIs together with agreement of both read-start positions. The maximum paired-UMI distance and start-coordinate tolerance were 1 and 3 bp, respectively; a retained duplex unit comprised two complementary paired-end fragments, corresponding to four primary alignments. The workflow did not generate strand-specific sequence consensuses and did not impose a minimum read-family size for SV calling.

### SV-associated breakpoint-junction calling and classification

A candidate SV-associated breakpoint junction was evaluated from one retained read pair from each complementary source strand. All four alignments were required to be primary, to map to an included chromosome and to have mapping quality ≥60. Secondary and supplementary alignments were rejected. The two complementary read pairs were required to show reciprocal mirrored mapping configurations with an overlap fraction >0.8, and the mates within a discordant pair were required not to overlap. A same-chromosome inward-facing configuration with an inferred interval of at least 1 kb was annotated as deletion-like (DEL), whereas a configuration connecting different chromosomes was annotated as translocation-like (TRN). Other same-chromosome orientations were recorded as duplication-like or inversion-like configurations, and same-chromosome deletion-like configurations below 1 kb were not included in the reported DEL counts. Junction records with the same class, chromosome pair and orientation and with both breakpoint coordinates separated by <500 bp were treated as matching, including an equivalent reversed representation. The measurement unit is therefore a duplex-supported breakpoint junction rather than a reconstructed complete structural variant.

### Breakpoint-frequency calculation and background correction

For each sample, the raw frequency for a reported junction class was calculated from the number of final somatic junctions divided by the cumulative length of qualifying proper duplex fragments. For each proper duplex fragment, the denominator contribution was the larger absolute template length recorded for its two complementary read pairs. Discordant duplex fragments did not contribute to this denominator. Raw frequencies were converted to events per 10⁹ qualifying duplex bases. For each bleomycin and irradiation sample, background-corrected deletion-like and translocation-like frequencies were calculated before group summaries by subtracting 0.15 and 2.85 per 10⁹ bases, respectively.

### Human–Drosophila species-mixing calibration

Human and Drosophila melanogaster genomic DNA were mixed at a 1:1 molar ratio and processed using the SMM-SV-seq library protocol. Five independently prepared mixed-DNA libraries were included. This composite reference contained the GRCh38 human primary chromosomes 1–22, X and Y; D. melanogaster BDGP6.46 contigs 2L, 2R, 3L, 3R, 4, X and Y; and Caenorhabditis elegans chromosomes I–V and X. No C. elegans DNA was included in the mixture. A junction with one end assigned to the human component and the other to the Drosophila component was counted as a cross-species artifact. The cross-species artifact rate for each library was calculated as the cross-species count divided by the cumulative qualifying proper-duplex length and expressed per 10⁹ bases.

### GM24385 annotated-deletion comparison

SMM-SV-seq deletion calls from GM24385 (HG002/NA24385) were compared with records in HG002_SVs_Tier1_v0.6.vcf.gz (GRCh37; file date 5 June 2018). Autosomal records annotated as homozygous deletions with an absolute SV length of at least 1 kb were selected. A deletion call was preferentially matched when both breakpoints were within 500 bp of the reference breakpoints; when no two-breakpoint match was available, a call with one breakpoint within this tolerance was accepted as a one-breakpoint match. Reference records were classified as reported, detected but not reported, covered without a breakpoint call, or not covered.

### Statistical analysis

Statistical analyses were performed using GraphPad Prism 11.1.0. Deletion-like and translocation-like breakpoint-junction frequencies were analyzed separately. For the bleomycin experiment, differences among groups were tested by ordinary one-way ANOVA followed by two-sided Dunnett tests comparing each bleomycin dose with the untreated group. For the irradiation experiment, breakpoint-junction frequencies were analyzed by two-way repeated-measures ANOVA, with genotype as the between-experiment factor and irradiation condition (0 or 2 Gy) as the within-experiment matched factor. The genotype-by-irradiation interaction tested whether the irradiation-associated change differed between the two cell lines. Prespecified within-genotype comparisons were evaluated using two-sided tests. Data are presented as mean ± s.d.; n=3 biological culture samples were analyzed per condition in each treatment experiment, and n=5 independently prepared libraries were used for the mixed-species calibration.

### Declaration of generative AI and AI-assisted technologies in the writing process

During the preparation of this work the author(s) used ChatGPT 5.6 in order to improve language and readability. After using this tool/service, the authors reviewed and edited the content as needed and take full responsibility for the content of the publication.

## Notes

### Competing Interest Statement

A.Y.M. and J.V. are co-founders of Mutagentech Corp.

### Summary of Updates

Title and abstract revised; GM24385 benchmarking updated with revised precision and recall calculations, caller comparisons, and breakpoint-coordinate analysis; bleomycin results expanded to include deletion-like junction spans; irradiation results and statistical analyses updated; Methods expanded to clarify library preparation, computational processing, filtering, and background correction; main figures redesigned and supplementary figures and tables updated; Discussion revised to clarify comparisons with previous methods and the scope of the assay.

