## Supplementary Figures and Tables for "Chimera-suppressing error-corrected sequencing enables quantitative detection of rare somatic structural-variant breakpoints in human cells"

#### Supplementary Figure S1.

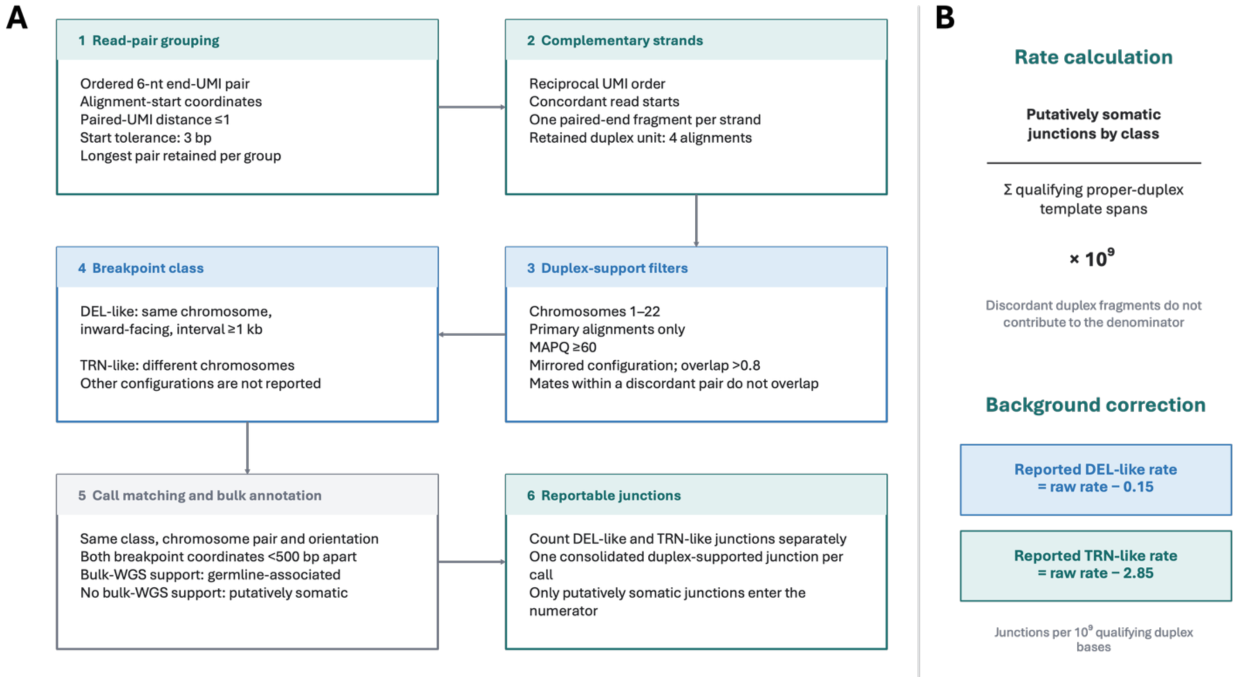

**Supplementary Figure S1. SMM-SV-seq computational workflow and breakpoint-frequency calculation.** (**A**) Overview of read-pair grouping, complementary-strand identification, duplex-support filtering, breakpoint-class assignment, call matching, and annotation using designated control data and matched conventional WGS. Each retained duplex unit comprised one paired-end fragment from each complementary source strand. Candidate junctions required four primary autosomal alignments with mapping quality ≥60 and mirrored support from the complementary strands. Same-chromosome inward-facing configurations spanning ≥1 kb were classified as deletion-like, whereas interchromosomal configurations were classified as translocation-like. Analysis-specific UMI, start-coordinate, and breakpoint-matching thresholds are provided in Methods. Junctions lacking evidence in the designated controls and other project libraries were classified as putatively somatic. (**B**) Raw class-specific frequency was calculated as the number of putatively somatic junctions divided by the cumulative template-span length of qualifying proper duplex fragments and expressed per $10^9$ bases. Discordant duplex fragments were excluded from the denominator. For the bleomycin and irradiation analyses, background-corrected rates were calculated by subtracting 0.15 deletion-like or 2.85 translocation-like junctions per $10^9$ qualifying duplex bases.

**Supplementary Figure S2.**

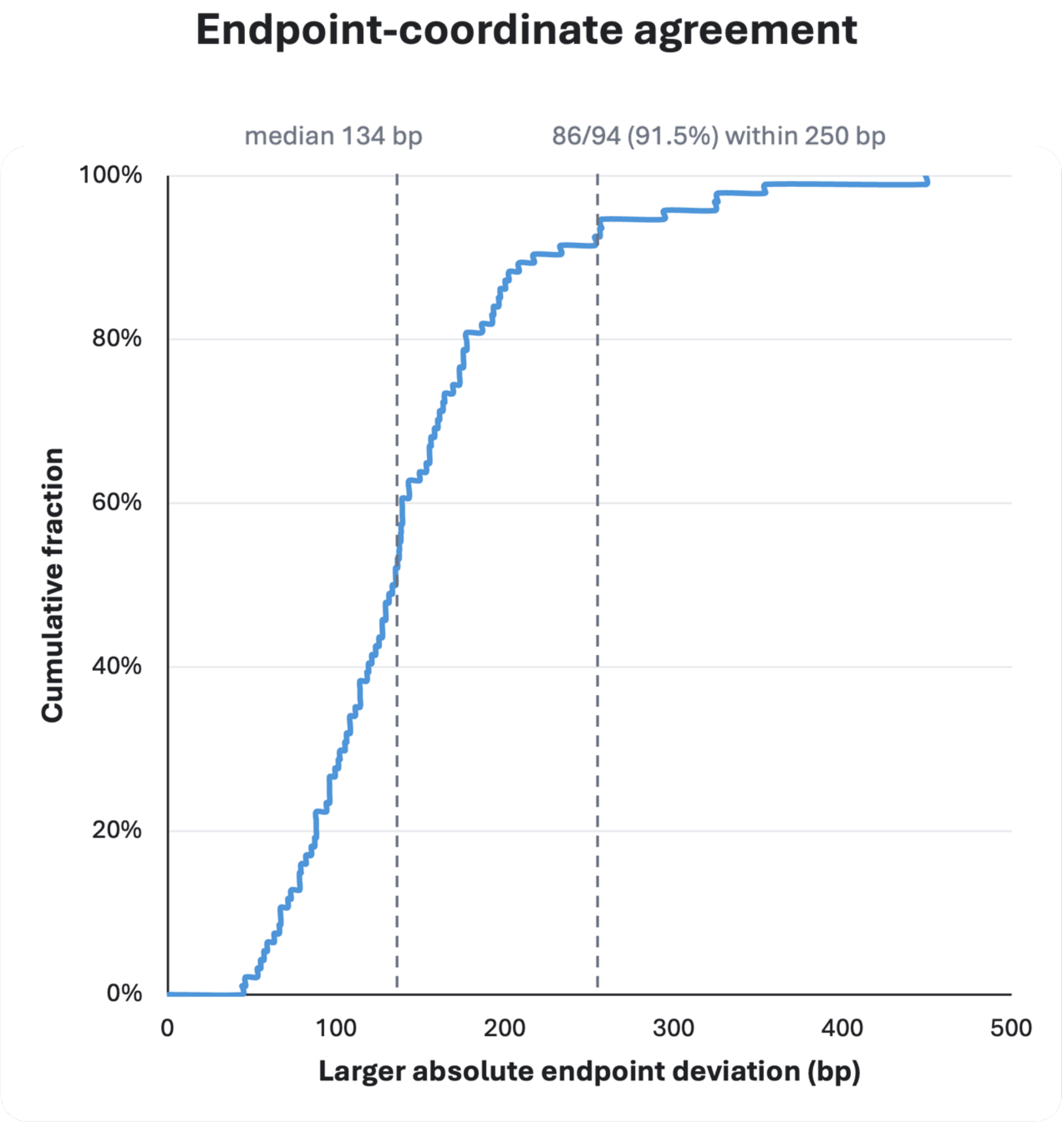

**Supplementary Figure S2. Endpoint-coordinate agreement in the GM24385 deletion benchmark.** Empirical cumulative distribution of the larger absolute deviation between the SMM-SV-seq and reference coordinates at the two deletion endpoints for the 94 calls matched at both breakpoints. The median larger endpoint deviation was 134 bp (interquartile range, 96–172 bp), and both endpoints were within 250 bp for 86 of 94 calls (91.5%). Calls were matched using a tolerance of 500 bp per endpoint.

### Supplementary Table 1.

| known_id | chrom | start | end | size_bp | zygosity | SMM_match | SMM_type | SMM_SAO | reason_not_match | SMM_chrom | SMM_start | SMM_end | SMM_size_bp | SMM_detection_type |
|---|---|---|---|---|---|---|---|---|---|---|---|---|---|---|
| HG3_Ill_GATKHCSBGrefine_157 | 1 | 25,158,687 | 25,161,509 | 2822 | hom | no | | | no_coverage | | | | | |
| HG3_PB_SVrefine2Falcon1Dovetail_293 | 1 | 53,594,100 | 53,595,603 | 1503 | hom | no | | | no_coverage | | | | | |
| HG3_Ill_SVrefine2DISCOVARDovetail_450 | 1 | 83,125,958 | 83,127,569 | 1611 | hom | yes | DEL | 1 | | 1 | 83125955 | 83127923 | 1968 | both_breakpoints |
| HG2_PB_SVrefine2Falcon1plusDovetail_434 | 1 | 84,517,925 | 84,524,629 | 6704 | hom | yes | DEL | 1 | | 1 | 84517776 | 84524754 | 6978 | both_breakpoints |
| HG2_PB_assemblyticsPBcR_1056 | 1 | 108,733,326 | 108,737,251 | 3925 | hom | yes | DEL | 1 | | 1 | 108733190 | 108737338 | 4148 | both_breakpoints |
| HG3_PB_SVrefine2PBcRDovetail_445 | 1 | 112,835,082 | 112,837,660 | 2578 | hom | no | | | low_MapQ | | | | | |
| HG4_PB_SVrefine2Falcon1Dovetail_552 | 1 | 156,526,704 | 156,528,935 | 2231 | hom | yes | DEL | 1 | | 1 | 156526590 | 156529033 | 2443 | both_breakpoints |
| HG3_PB_SVrefine2PBcRDovetail_565 | 1 | 184,814,728 | 184,820,784 | 6056 | hom | yes | DEL | 1 | | 1 | 184814601 | 184820812 | 6211 | both_breakpoints |
| HG4_PB_SVrefine2Falcon1Dovetail_681 | 1 | 197,500,508 | 197,503,271 | 2763 | hom | no | | | no_coverage | | | | | |
| HG4_PB_SVrefine2Falcon1Dovetail_712 | 1 | 207,541,963 | 207,545,564 | 3601 | hom | yes | DEL | 1 | | 1 | 207541956 | 207545687 | 3731 | both_breakpoints |
| HG4_Ill_GATKHCSBGrefine_772 | 1 | 212,471,135 | 212,472,616 | 1481 | hom | yes | DEL | 1 | | 1 | 212471034 | 212472678 | 1644 | both_breakpoints |
| HG2_10X_SVrefine210Xhap12_889 | 1 | 213,084,227 | 213,086,169 | 1942 | hom | no | | | low_MapQ | | | | | |
| HG2_10X_SVrefine210Xhap12_904 | 1 | 218,182,488 | 218,188,594 | 6106 | hom | no | | | no_coverage | | | | | |
| HG2_PB_SVrefine2Falcon1plusDovetail_872 | 1 | 233,962,209 | 233,963,552 | 1343 | hom | no | | | no_coverage | | | | | |
| HG3_PB_SVrefine2PBcRDovetail_755 | 1 | 246,137,767 | 246,139,255 | 1488 | hom | no | | | no_coverage | | | | | |
| HG4_PB_SVrefine2Falcon1Dovetail_917 | 1 | 247,850,455 | 247,856,508 | 6053 | hom | no | | | no_coverage | | | | | |
| HG4_PB_SVrefine2Falcon1Dovetail_919 | 1 | 248,051,493 | 248,057,699 | 6206 | hom | yes | DEL | 1 | | 1 | 248051307 | 248057639 | 6332 | both_breakpoints |
| HG3_Ill_SVrefine2DISCOVARDovetail_1082 | 2 | 4,781,283 | 4,787,348 | 6065 | hom | yes | DEL | 1 | | 2 | 4781164 | 4787357 | 6193 | both_breakpoints |
| HG2_10X_SVrefine210Xhap12_1252 | 2 | 9,925,472 | 9,929,048 | 3576 | hom | no | | | no_coverage | | | | | |
| HG2_PB_SVrefine2Falcon2Bionano_619 | 2 | 16,271,567 | 16,273,799 | 2232 | hom | no | | | no_coverage | | | | | |
| HG2_PB_assemblyticsPBcR_12878 | 2 | 31,725,177 | 31,727,388 | 2211 | hom | no | | | no_coverage | | | | | |
| HG2_PB_HySA_3330 | 2 | 33,141,282 | 33,143,464 | 2182 | hom | no | | | no_coverage | | | | | |
| HG2_PB_SVrefine2PBcRplusDovetail_5587 | 2 | 36,216,159 | 36,218,225 | 2066 | hom | no | | | no_SV_call | | | | | |
| HG2_Ill_SVrefine2DISCOVARplusDovetail_6494 | 2 | 41,775,967 | 41,781,283 | 5316 | hom | yes | DEL | 1 | | 2 | 41775896 | 41781351 | 5455 | both_breakpoints |
| HG2_PB_SVrefine2Falcon2Bionano_693 | 2 | 41,973,141 | 41,975,865 | 2724 | hom | no | | | no_coverage | | | | | |
| HG4_Ill_SVrefine2DISCOVARDovetail_1286 | 2 | 42,346,260 | 42,347,318 | 1058 | hom | no | | | no_coverage | | | | | |
| HG2_10X_SVrefine210Xhap12_1517 | 2 | 76,526,498 | 76,528,893 | 2395 | hom | yes | DEL | 1 | | 2 | 76526363 | 76528968 | 2605 | both_breakpoints |
| HG2_PB_SVrefine2Falcon1plusDovetail_6156 | 2 | 76,773,554 | 76,775,452 | 1898 | hom | no | | | no_coverage | | | | | |
| HG2_Ill_MetaSV_2548 | 2 | 89,029,228 | 89,032,282 | 3054 | hom | no | | | no_coverage | | | | | |
| HG3_PB_pbsv_2100 | 2 | 105,825,891 | 105,828,426 | 2535 | hom | no | | | no_coverage | | | | | |
| HG3_PB_pbsv_2227 | 2 | 126,443,247 | 126,451,831 | 8584 | hom | yes | DEL | 1 | | 2 | 126443110 | 126451833 | 8723 | both_breakpoints |
| HG2_PB_pbsv_2252 | 2 | 127,674,756 | 127,677,272 | 2516 | hom | no | | | no_coverage | | | | | |
| HG2_PB_SVrefine2PB10Xhap12_1818 | 2 | 159,958,799 | 159,961,068 | 2269 | hom | no | | | no_SV_call | | | | | |
| HG4_PB_assemblyticsfalcon_14061 | 2 | 160,140,080 | 160,158,998 | 18918 | hom | no | | | no_coverage | | | | | |
| HG3_PB_SVrefine2PBcRDovetail_1218 | 2 | 160,149,117 | 160,151,648 | 2531 | hom | yes | DEL | 1 | | 2 | 160149039 | 160151714 | 2675 | both_breakpoints |
| HG4_Ill_SVrefine2DISCOVARDovetail_1791 | 2 | 162,332,267 | 162,334,041 | 1774 | hom | yes | DEL | 1 | | 2 | 162332050 | 162334042 | 1992 | both_breakpoints |
| HG4_Ill_GATKHCSBGrefine_1690 | 2 | 170,641,507 | 170,643,951 | 2444 | hom | no | | | no_coverage | | | | | |
| HG3_PB_SVrefine2Falcon1Dovetail_1615 | 2 | 202,146,648 | 202,149,440 | 2792 | hom | yes | DEL | 1 | | 2 | 202146645 | 202149536 | 2891 | both_breakpoints |
| HG2_PB_SVrefine2PBcRplusDovetail_6108 | 2 | 208,474,478 | 208,476,780 | 2302 | hom | no | | | no_coverage | | | | | |
| HG2_PB_SVrefine2PBcRplusDovetail_6154 | 2 | 223,760,178 | 223,762,671 | 2493 | hom | yes | DEL | 1 | | 2 | 223759986 | 223762685 | 2699 | both_breakpoints |
| HG2_PB_SVrefine2Falcon2Bionano_1175 | 2 | 227,334,127 | 227,335,281 | 1154 | hom | no | | | no_coverage | | | | | |
| HG2_PB_SVrefine2PB10Xhap12_2033 | 2 | 229,264,344 | 229,268,336 | 3992 | hom | yes | DEL | 1 | | 2 | 229264297 | 229268497 | 4200 | both_breakpoints |
| HG2_PB_assemblyticsfalcon_14974 | 2 | 235,546,753 | 235,556,629 | 9876 | hom | no | | | no_SV_call | | | | | |
| HG3_PB_SVrefine2PBcRDovetail_1458 | 2 | 242,704,671 | 242,706,171 | 1500 | hom | no | | | no_coverage | | | | | |
| HG4_Ill_GATKHCSBGrefine_2063 | 3 | 4,066,924 | 4,069,555 | 2631 | hom | yes | DEL | 1 | | 3 | 4066917 | 4069692 | 2775 | both_breakpoints |
| HG2_PB_SVrefine2PBcRplusDovetail_7299 | 3 | 25,030,888 | 25,032,249 | 1361 | hom | yes | DEL | 1 | | 3 | 25030887 | 25032412 | 1525 | both_breakpoints |
| HG2_PB_SVrefine2Falcon2Bionano_1402 | 3 | 26,450,969 | 26,452,298 | 1329 | hom | no | | | no_coverage | | | | | |
| HG2_PB_SVrefine2Falcon1plusDovetail_7896 | 3 | 32,102,050 | 32,107,883 | 5833 | hom | yes | DEL | 1 | | 3 | 32101963 | 32107899 | 5936 | both_breakpoints |
| HG3_Ill_SVrefine2DISCOVARDovetail_2340 | 3 | 32,806,536 | 32,807,843 | 1307 | hom | no | | | no_coverage | | | | | |
| HG4_Ill_GATKHCSBGrefine_2235 | 3 | 42,838,356 | 42,840,810 | 2454 | hom | yes | DEL | 1 | | 3 | 42838148 | 42840821 | 2673 | both_breakpoints |
| HG2_10X_SVrefine210Xhap12_2512 | 3 | 67,493,695 | 67,496,799 | 3104 | hom | no | | | no_coverage | | | | | |
| HG2_10X_SVrefine210Xhap12_2515 | 3 | 68,739,683 | 68,747,858 | 8175 | hom | yes | DEL | 1 | | 3 | 68739648 | 68747972 | 8324 | both_breakpoints |
| HG2_PB_SVrefine2Falcon2Bionano_1571 | 3 | 80,925,311 | 80,926,604 | 1293 | hom | yes | DEL | 1 | | 3 | 80925310 | 80926759 | 1449 | both_breakpoints |
| HG4_PB_SVrefine2Falcon1Dovetail_2080 | 3 | 83,853,115 | 83,855,349 | 2234 | hom | no | | | no_coverage | | | | | |
| HG2_PB_SVrefine2PB10Xhap12_2556 | 3 | 98,410,705 | 98,414,783 | 4078 | hom | no | | | no_coverage | | | | | |
| HG4_PB_SVrefine2Falcon1Dovetail_2112 | 3 | 98,899,060 | 98,902,389 | 3329 | hom | yes | DEL | 1 | | 3 | 98898883 | 98902390 | 3507 | both_breakpoints |
| HG2_PB_SVrefine2Falcon2Bionano_1736 | 3 | 146,385,191 | 146,390,404 | 5213 | hom | yes | DEL | 1 | | 3 | 146385047 | 146394874 | 9827 | one_breakpoint |
| HG2_PB_SVrefine2Falcon1plusDovetail_8262 | 3 | 152,311,723 | 152,313,155 | 1432 | hom | yes | DEL | 1 | | 3 | 152311594 | 152313156 | 1562 | both_breakpoints |
| HG3_Ill_SVrefine2DISCOVARDovetail_2946 | 3 | 162,764,902 | 162,769,084 | 4182 | hom | yes | DEL | 1 | | 3 | 162764742 | 162769118 | 4376 | both_breakpoints |
| HG3_PB_SVrefine2PBcRDovetail_1847 | 3 | 186,581,015 | 186,585,284 | 4269 | hom | yes | DEL | 1 | | 3 | 186580965 | 186585405 | 4440 | both_breakpoints |

| | | | | | | | | | | | | | | |
|---|---|---|---|---|---|---|---|---|---|---|---|---|---|---|
| HG3_PB_SVrefine2PBcRDovetail_1970 | 4 | 6,651,589 | 6,652,910 | 1321 | hom | no | | | no_coverage | | | | | |
| HG3_PB_SVrefine2Falcon1Dovetail_2673 | 4 | 10,211,260 | 10,234,567 | 23307 | hom | yes | DEL | 1 | | 4 | 10211122 | 10234675 | 23553 | both_breakpoints |
| HG4_Ill_SVrefine2DISCOVARDovetail_3360 | 4 | 40,024,907 | 40,026,274 | 1367 | hom | no | | | no_coverage | | | | | |
| HG4_Ill_GATKHCSBGrefine_3121 | 4 | 46,202,795 | 46,205,026 | 2231 | hom | no | | | no_coverage | | | | | |
| HG3_PB_assemblyticsfalcon_19575 | 4 | 75,642,742 | 75,648,799 | 6057 | hom | no | | | low_MapQ | | | | | |
| HG4_Ill_MetaSV_3788 | 4 | 79,269,120 | 79,275,195 | 6075 | hom | no | | | no_coverage | | | | | |
| HG4_Ill_SVrefine2DISCOVARDovetail_3596 | 4 | 91,596,785 | 91,602,908 | 6123 | hom | yes | DEL | 1 | | 4 | 91596798 | 91603083 | 6285 | both_breakpoints |
| HG4_PB_SVrefine2PBcRDovetail_2410 | 4 | 92,159,979 | 92,161,231 | 1252 | hom | no | | | no_SV_call | | | | | |
| HG3_PB_pbsv_4664 | 4 | 108,127,808 | 108,131,714 | 3906 | hom | yes | DEL | 1 | | 4 | 108127779 | 108131887 | 4108 | both_breakpoints |
| HG2_PB_SVrefine2PBcRplusDovetail_8363 | 4 | 135,432,950 | 135,435,701 | 2751 | hom | no | | | no_coverage | | | | | |
| HG2_PB_SVrefine2PBcRplusDovetail_8371 | 4 | 139,468,835 | 139,473,221 | 4386 | hom | yes | DEL | 1 | | 4 | 139468797 | 139473267 | 4470 | both_breakpoints |
| HG4_Ill_MetaSV_3913 | 4 | 142,230,778 | 142,233,062 | 2284 | hom | yes | DEL | 1 | | 4 | 142230684 | 142233063 | 2379 | both_breakpoints |
| HG2_Ill_GATKHCSBGrefine_3395 | 4 | 146,438,806 | 146,440,450 | 1644 | hom | yes | DEL | 1 | | 4 | 146438668 | 146440473 | 1805 | both_breakpoints |
| HG4_PB_SVrefine2Falcon1Dovetail_3071 | 4 | 159,393,249 | 159,394,601 | 1352 | hom | no | | | no_SV_call | | | | | |
| HG2_10X_SVrefine210Xhap12_3708 | 4 | 161,879,029 | 161,884,943 | 5914 | hom | yes | DEL | 1 | | 4 | 161878989 | 161885009 | 6020 | both_breakpoints |
| HG4_PB_SVrefine2Falcon1Dovetail_3104 | 4 | 167,677,025 | 167,683,060 | 6035 | hom | yes | DEL | 1 | | 4 | 167676930 | 167683189 | 6259 | both_breakpoints |
| HG2_PB_assemblyticsPBcR_19771 | 4 | 172,988,640 | 172,992,931 | 4291 | hom | no | | | no_coverage | | | | | |
| HG4_Ill_SVrefine2DISCOVARDovetail_3964 | 4 | 181,876,786 | 181,878,288 | 1502 | hom | yes | DEL | 1 | | 4 | 181876647 | 181878360 | 1713 | both_breakpoints |
| HG3_PB_SVrefine2PBcRDovetail_2430 | 4 | 186,441,635 | 186,444,073 | 2438 | hom | yes | DEL | 1 | | 4 | 186441539 | 186444091 | 2552 | both_breakpoints |
| HG2_PB_SVrefine2PBcRplusDovetail_8739 | 5 | 3,637,249 | 3,638,404 | 1155 | hom | no | | | no_coverage | | | | | |
| HG2_Ill_MetaSV_3926 | 5 | 24,370,527 | 24,373,436 | 2909 | hom | no | | | no_coverage | | | | | |
| HG3_PB_SVrefine2PBcRDovetail_2654 | 5 | 42,165,928 | 42,167,769 | 1841 | hom | yes | DEL | 1 | | 5 | 42165860 | 42167842 | 1982 | both_breakpoints |
| HG2_PB_pbsv_5780 | 5 | 46,270,652 | 46,275,836 | 5184 | hom | yes | DEL | 1 | | 5 | 46270635 | 46276032 | 5397 | both_breakpoints |
| HG3_Ill_SVrefine2DISCOVARDovetail_4325 | 5 | 55,428,997 | 55,430,134 | 1137 | hom | yes | DEL | 1 | | 5 | 55429156 | 55430428 | 1272 | both_breakpoints |
| HG3_Ill_SVrefine2DISCOVARDovetail_4337 | 5 | 57,679,986 | 57,686,095 | 6109 | hom | yes | DEL | 1 | | 5 | 57679890 | 57686122 | 6232 | both_breakpoints |
| HG2_Ill_SpiralSDKrefine_13579 | 5 | 60,001,704 | 60,003,665 | 1961 | hom | no | | | no_coverage | | | | | |
| HG3_Ill_SVrefine2DISCOVARDovetail_4439 | 5 | 83,947,917 | 83,954,879 | 6962 | hom | no | | | no_SV_call | | | | | |
| HG2_PB_SVrefine2PB10Xhap12_4329 | 5 | 90,499,772 | 90,502,095 | 2323 | hom | yes | DEL | 1 | | 5 | 90499614 | 90502118 | 2504 | both_breakpoints |
| HG2_PB_SVrefine2PB10Xhap12_4349 | 5 | 98,345,118 | 98,347,482 | 2364 | hom | yes | DEL | 1 | | 5 | 98345117 | 98347549 | 2432 | both_breakpoints |
| HG4_Ill_SVrefine2DISCOVARDovetail_4513 | 5 | 119,380,147 | 119,382,680 | 2533 | hom | no | | | no_coverage | | | | | |
| HG2_PB_SVrefine2PBcRplusDovetail_9127 | 5 | 136,854,637 | 136,856,353 | 1716 | hom | no | | | no_coverage | | | | | |
| HG2_PB_assemblyticsfalcon_22079 | 5 | 147,553,040 | 147,554,778 | 1738 | hom | no | | | no_coverage | | | | | |
| HG4_Ill_MetaSV_4307 | 5 | 151,456,413 | 151,462,447 | 6034 | hom | yes | DEL | 1 | | 5 | 151456314 | 151462529 | 6215 | both_breakpoints |
| HG2_PB_HySA_11901 | 5 | 176,387,579 | 176,390,180 | 2601 | hom | yes | DEL | 1 | | 5 | 176387424 | 176390218 | 2794 | both_breakpoints |
| HG2_Ill_SVrefine2DISCOVARplusDovetail_10865 | 6 | 666,376 | 667,821 | 1445 | hom | yes | DEL | 1 | | 6 | 666272 | 667929 | 1657 | both_breakpoints |
| HG2_Ill_SVrefine2DISCOVARplusDovetail_10896 | 6 | 5,037,250 | 5,039,186 | 1936 | hom | yes | DEL | 1 | | 6 | 5037194 | 5039292 | 2098 | both_breakpoints |
| HG4_Ill_GATKHCSBGrefine_4595 | 6 | 5,037,643 | 5,039,186 | 1543 | hom | yes | DEL | 1 | | 6 | 5037194 | 5039292 | 2098 | both_breakpoints |
| HG3_Ill_SVrefine2DISCOVARDovetail_5066 | 6 | 24,811,875 | 24,817,950 | 6075 | hom | yes | DEL | 1 | | 6 | 24811877 | 24818035 | 6158 | both_breakpoints |
| HG3_Ill_GATKHCSBGrefine_4587 | 6 | 29,685,469 | 29,688,080 | 2611 | hom | yes | DEL | 1 | | 6 | 29685388 | 29688211 | 2823 | both_breakpoints |
| HG2_Ill_GATKHCSBGrefine_4658 | 6 | 29,899,768 | 29,901,526 | 1758 | hom | no | | | low_MapQ | | | | | |
| HG2_Ill_MetaSV_4288 | 6 | 32,499,303 | 32,500,806 | 1503 | hom | no | | | no_coverage | | | | | |
| HG3_Ill_MetaSV_4465 | 6 | 32,561,955 | 32,564,608 | 2653 | hom | yes | DEL | 1 | | 6 | 32561837 | 32564656 | 2819 | both_breakpoints |
| HG4_PB_SVrefine2Falcon1Dovetail_4158 | 6 | 53,928,710 | 53,934,828 | 6118 | hom | no | | | no_coverage | | | | | |
| HG2_PB_SVrefine2Falcon2Bionano_3489 | 6 | 66,260,027 | 66,262,021 | 1994 | hom | no | | | no_coverage | | | | | |
| HG2_PB_SVrefine2PB10Xhap12_5129 | 6 | 74,865,433 | 74,866,584 | 1151 | hom | no | | | no_coverage | | | | | |
| HG3_PB_SVrefine2Falcon1Dovetail_4245 | 6 | 77,097,492 | 77,102,641 | 5149 | hom | yes | DEL | 1 | | 6 | 77097319 | 77102656 | 5337 | both_breakpoints |
| HG3_PB_SVrefine2Falcon1Dovetail_4272 | 6 | 85,318,143 | 85,324,220 | 6077 | hom | yes | DEL | 1 | | 6 | 85318058 | 85324413 | 6355 | both_breakpoints |
| HG4_Ill_GATKHCSBGrefine_5002 | 6 | 86,708,739 | 86,714,791 | 6052 | hom | no | | | low_MapQ | | | | | |
| HG4_Ill_SVrefine2DISCOVARDovetail_5271 | 6 | 95,193,322 | 95,194,336 | 1014 | hom | yes | DEL | 1 | | 6 | 95193265 | 95194384 | 1119 | both_breakpoints |
| HG3_PB_SVrefine2PBcRDovetail_3321 | 6 | 108,031,343 | 108,032,402 | 1059 | hom | no | | | no_coverage | | | | | |
| HG3_Ill_MetaSV_4647 | 6 | 129,319,515 | 129,325,574 | 6059 | hom | no | | | no_coverage | | | | | |
| HG3_PB_SVrefine2PBcRDovetail_3331 | 6 | 133,341,820 | 133,347,885 | 6065 | hom | no | | | no_coverage | | | | | |
| HG2_PB_SVrefine2PBcRplusDovetail_9809 | 6 | 151,485,839 | 151,486,912 | 1073 | hom | no | | | no_coverage | | | | | |
| HG3_Ill_SVrefine2DISCOVARDovetail_5794 | 6 | 168,716,918 | 168,718,524 | 1606 | hom | no | | | no_coverage | | | | | |
| HG2_PB_SVrefine2PBcRplusDovetail_9999 | 7 | 224,004 | 225,192 | 1188 | hom | no | | | no_coverage | | | | | |
| HG4_10X_allpass_1554 | 7 | 24,038,161 | 24,040,070 | 1909 | hom | yes | DEL | 1 | | 7 | 24038060 | 24040213 | 2153 | both_breakpoints |
| HG2_Ill_breakscan11_1168 | 7 | 32,390,151 | 32,393,112 | 2961 | hom | no | | | no_coverage | | | | | |
| HG4_PB_SVrefine2PBcRDovetail_4012 | 7 | 49,719,838 | 49,725,897 | 6059 | hom | no | | | no_coverage | | | | | |
| HG2_PB_SVrefine2Falcon1plusDovetail_11086 | 7 | 64,315,807 | 64,318,058 | 2251 | hom | yes | DEL | 1 | | 7 | 64315554 | 64318059 | 2505 | both_breakpoints |
| HG3_10X_allpass_1670 | 7 | 73,549,758 | 73,550,966 | 1208 | hom | yes | DEL | 1 | | 7 | 73549711 | 73551166 | 1455 | both_breakpoints |
| HG2_PB_SVrefine2PB10Xhap12_6030 | 7 | 73,828,443 | 73,831,261 | 2818 | hom | no | | | no_coverage | | | | | |
| HG2_PB_SVrefine2PB10Xhap12_6107 | 7 | 89,810,407 | 89,812,599 | 2192 | hom | no | | | low_MapQ | | | | | |
| HG4_Ill_SVrefine2DISCOVARDovetail_6128 | 7 | 96,475,886 | 96,481,994 | 6108 | hom | yes | DEL | 1 | | 7 | 96475879 | 96482171 | 6292 | both_breakpoints |
| HG2_PB_PB10Xdip_26327 | 7 | 109,436,689 | 109,453,905 | 17216 | hom | no | | | no_coverage | | | | | |
| HG2_Ill_MetaSV_4825 | 7 | 113,416,161 | 113,422,207 | 6046 | hom | yes | DEL | 1 | | 7 | 113416036 | 113422215 | 6179 | both_breakpoints |

| | | | | | | | | | | | | | | |
|---|---|---|---|---|---|---|---|---|---|---|---|---|---|---|
| HG3_Ill_SVrefine2DISCOVARDovetail_6533 | 7 | 148,072,863 | 148,076,323 | 3460 | hom | yes | DEL | 1 | | 7 | 148072850 | 148076428 | 3578 | both_breakpoints |
| HG2_PB_SVrefine2Falcon1plusDovetail_11702 | 8 | 25,066,678 | 25,070,652 | 3974 | hom | yes | DEL | 1 | | 8 | 25066599 | 25070699 | 4100 | both_breakpoints |
| HG4_PB_pbsv_9369 | 8 | 32,680,005 | 32,691,261 | 11256 | hom | no | | | no_coverage | | | | | |
| HG3_PB_assemblyticsfalcon_26302 | 8 | 42,185,330 | 42,194,208 | 8878 | hom | yes | DEL | 1 | | 8 | 42190388 | 42194532 | 4144 | one_breakpoint |
| HG2_10X_SVrefine210Xhap12_6668 | 8 | 42,190,248 | 42,194,208 | 3960 | hom | yes | DEL | 1 | | 8 | 42190388 | 42194532 | 4144 | both_breakpoints |
| HG2_PB_assemblyticsfalcon_26629 | 8 | 75,362,872 | 75,366,976 | 4104 | hom | no | | | no_coverage | | | | | |
| HG3_PB_SVrefine2Falcon1Dovetail_5830 | 8 | 87,188,248 | 87,193,743 | 5495 | hom | no | | | no_coverage | | | | | |
| HG2_PB_SVrefine2Falcon1plusDovetail_12022 | 8 | 126,595,121 | 126,601,134 | 6013 | hom | no | | | no_coverage | | | | | |
| HG2_Ill_GATKHCSBGrefine_6676 | 8 | 127,192,207 | 127,194,571 | 2364 | hom | no | | | no_coverage | | | | | |
| HG3_PB_SVrefine2Falcon1Dovetail_5976 | 8 | 129,465,153 | 129,471,266 | 6113 | hom | no | | | no_coverage | | | | | |
| HG3_PB_SVrefine2PBcRDovetail_4488 | 8 | 135,082,913 | 135,089,015 | 6102 | hom | yes | DEL | 1 | | 8 | 135082869 | 135089034 | 6165 | both_breakpoints |
| HG4_PB_HySA_14406 | 8 | 142,643,076 | 142,644,203 | 1127 | hom | no | | | no_SV_call | | | | | |
| HG3_Ill_SVrefine2DISCOVARDovetail_7485 | 9 | 10,020,266 | 10,021,396 | 1130 | hom | no | | | no_SV_call | | | | | |
| HG2_PB_SVrefine2Falcon2Bionano_4789 | 9 | 17,908,785 | 17,912,150 | 3365 | hom | no | | | no_SV_call | | | | | |
| HG2_PB_SVrefine2PBcRplusDovetail_11480 | 9 | 22,496,528 | 22,504,344 | 7816 | hom | no | | | low_MapQ | | | | | |
| HG4_10X_allpass_1977 | 9 | 31,291,355 | 31,292,672 | 1317 | hom | yes | DEL | 1 | | 9 | 31291212 | 31292778 | 1566 | both_breakpoints |
| HG2_PB_SVrefine2Falcon1plusDovetail_12308 | 9 | 33,423,361 | 33,424,654 | 1293 | hom | no | | | no_coverage | | | | | |
| HG4_PB_SVrefine2Falcon1Dovetail_6345 | 9 | 84,324,348 | 84,326,926 | 2578 | hom | no | | | low_MapQ | | | | | |
| HG4_PB_SVrefine2Falcon1Dovetail_6354 | 9 | 88,722,221 | 88,723,236 | 1015 | hom | no | | | no_SV_call | | | | | |
| HG2_PB_SVrefine2Falcon1plusDovetail_12422 | 9 | 89,154,558 | 89,155,946 | 1388 | hom | yes | DEL | 1 | | 9 | 89154499 | 89155947 | 1448 | both_breakpoints |
| HG2_Ill_SVrefine2DISCOVARplusDovetail_13642 | 9 | 90,858,983 | 90,860,963 | 1980 | hom | no | | | no_coverage | | | | | |
| HG2_Ill_GATKHCSBGrefine_7137 | 9 | 101,309,043 | 101,311,664 | 2621 | hom | yes | DEL | 1 | | 9 | 101308941 | 101311744 | 2803 | both_breakpoints |
| HG3_PB_SVrefine2Falcon1Dovetail_6346 | 9 | 110,018,277 | 110,020,848 | 2571 | hom | yes | DEL | 1 | | 9 | 110018210 | 110020883 | 2673 | both_breakpoints |
| HG3_PB_SVrefine2Falcon1Dovetail_6347 | 9 | 110,033,441 | 110,035,444 | 2003 | hom | no | | | no_coverage | | | | | |
| HG3_PB_SVrefine2Falcon1Dovetail_6349 | 9 | 110,537,520 | 110,540,595 | 3075 | hom | yes | DEL | 1 | | 9 | 110537323 | 110540703 | 3380 | both_breakpoints |
| HG2_PB_SVrefine2PB10Xhap12_7754 | 9 | 132,026,685 | 132,028,069 | 1384 | hom | yes | DEL | 1 | | 9 | 132026529 | 132028099 | 1570 | both_breakpoints |
| HG2_PB_assemblyticsPBcR_27603 | 9 | 138,470,556 | 138,480,199 | 9643 | hom | no | | | no_SV_call | | | | | |
| HG3_Ill_SVrefine2DISCOVARDovetail_7972 | 9 | 138,479,100 | 138,480,199 | 1099 | hom | no | | | no_coverage | | | | | |
| HG2_PB_SVrefine2PBcRplusDovetail_11888 | 9 | 140,253,510 | 140,255,960 | 2450 | hom | no | | | no_coverage | | | | | |
| HG4_PB_SVrefine2PBcRDovetail_5537 | 10 | 6,411,563 | 6,417,629 | 6066 | hom | yes | DEL | 1 | | 10 | 6411510 | 6417653 | 6143 | both_breakpoints |
| HG2_Ill_SVrefine2DISCOVARplusDovetail_1304 | 10 | 24,376,838 | 24,378,441 | 1603 | hom | no | | | low_MapQ | | | | | |
| HG2_PB_SVrefine2Falcon2Bionano_5371 | 10 | 31,443,332 | 31,444,846 | 1514 | hom | no | | | no_coverage | | | | | |
| HG2_Ill_GATKHCSBGrefine_7714 | 10 | 78,255,572 | 78,261,019 | 5447 | hom | yes | DEL | 1 | | 10 | 78255509 | 78261050 | 5541 | both_breakpoints |
| HG3_PB_pbsv_11205 | 10 | 93,633,335 | 93,634,562 | 1227 | hom | no | | | no_coverage | | | | | |
| HG3_PB_SVrefine2Falcon1Dovetail_6930 | 10 | 94,134,600 | 94,137,653 | 3053 | hom | no | | | no_coverage | | | | | |
| HG3_PB_SVrefine2Falcon1Dovetail_6945 | 10 | 99,034,863 | 99,037,397 | 2534 | hom | no | | | no_coverage | | | | | |
| HG2_PB_SVrefine2PBcRplusDovetail_1383 | 10 | 108,030,317 | 108,032,541 | 2224 | hom | yes | DEL | 1 | | 10 | 108030289 | 108032629 | 2340 | both_breakpoints |
| HG2_Ill_MetaSV_659 | 10 | 111,572,110 | 111,578,216 | 6106 | hom | yes | DEL | 1 | | 10 | 111571975 | 111578237 | 6262 | both_breakpoints |
| HG4_PB_SVrefine2Falcon1Dovetail_7078 | 10 | 122,226,594 | 122,228,743 | 2149 | hom | no | | | low_MapQ | | | | | |
| HG2_10X_SVrefine210Xhap12_8436 | 11 | 7,134,664 | 7,136,443 | 1779 | hom | no | | | no_coverage | | | | | |
| HG2_PB_SVrefine2PBcRplusDovetail_1640 | 11 | 7,790,210 | 7,791,886 | 1676 | hom | no | | | low_MapQ | | | | | |
| HG2_10X_SVrefine210Xhap12_8446 | 11 | 9,323,570 | 9,324,580 | 1010 | hom | no | | | no_coverage | | | | | |
| HG2_PB_SVrefine2PBcRplusDovetail_1790 | 11 | 47,057,589 | 47,063,706 | 6117 | hom | yes | DEL | 1 | | 11 | 47057588 | 47063820 | 6232 | both_breakpoints |
| HG2_PB_SVrefine2PBcRplusDovetail_1833 | 11 | 57,818,103 | 57,819,103 | 1000 | hom | no | | | no_SV_call | | | | | |
| HG3_PB_SVrefine2PBcRDovetail_5808 | 11 | 66,711,470 | 66,713,275 | 1805 | hom | no | | | no_coverage | | | | | |
| HG4_Ill_MetaSV_890 | 11 | 93,154,136 | 93,160,197 | 6061 | hom | no | | | no_coverage | | | | | |
| HG3_Ill_MetaSV_941 | 11 | 93,679,229 | 93,681,423 | 2194 | hom | no | | | no_coverage | | | | | |
| HG2_PB_SVrefine2Falcon2Bionano_6384 | 12 | 9,016,840 | 9,017,867 | 1027 | hom | yes | DEL | 1 | | 12 | 9016834 | 9017955 | 1121 | both_breakpoints |
| HG2_PB_SVrefine2PBcRplusDovetail_2252 | 12 | 13,545,544 | 13,551,618 | 6074 | hom | no | | | no_coverage | | | | | |
| HG3_Ill_GATKHCSBGrefine_8588 | 12 | 17,444,825 | 17,445,937 | 1112 | hom | no | | | no_coverage | | | | | |
| HG4_PB_SVrefine2Falcon1Dovetail_8008 | 12 | 43,977,449 | 43,979,549 | 2100 | hom | yes | DEL | 1 | | 12 | 43977394 | 43979587 | 2193 | both_breakpoints |
| HG2_PB_SVrefine2PB10Xhap12_9512 | 12 | 48,708,574 | 48,710,703 | 2129 | hom | no | | | no_coverage | | | | | |
| HG2_PB_SVrefine2PBcRplusDovetail_2340 | 12 | 48,725,466 | 48,728,380 | 2914 | hom | yes | DEL | 1 | | 12 | 48725313 | 48728380 | 3067 | both_breakpoints |
| HG4_Ill_MetaSV_1135 | 12 | 71,602,357 | 71,604,473 | 2116 | hom | yes | DEL | 1 | | 12 | 71602337 | 71604606 | 2269 | both_breakpoints |
| HG3_PB_SVrefine2PBcRDovetail_6275 | 12 | 72,356,508 | 72,358,072 | 1564 | hom | no | | | no_coverage | | | | | |
| HG3_PB_SVrefine2Falcon1Dovetail_8108 | 12 | 96,233,581 | 96,236,327 | 2746 | hom | no | | | no_coverage | | | | | |
| HG2_PB_SVrefine2PB10Xhap12_9929 | 12 | 132,879,957 | 132,881,349 | 1392 | hom | no | | | no_coverage | | | | | |
| HG4_PB_SVrefine2PBcRDovetail_6925 | 13 | 27,050,041 | 27,052,186 | 2145 | hom | no | | | low_MapQ | | | | | |
| HG2_PB_SVrefine2PB10Xhap12_10144 | 13 | 49,951,395 | 49,953,856 | 2461 | hom | yes | DEL | 1 | | 13 | 49951367 | 49953938 | 2571 | both_breakpoints |
| HG2_10X_allpass_2249 | 13 | 61,286,423 | 61,287,488 | 1065 | hom | no | | | no_coverage | | | | | |
| HG3_Ill_SVrefine2DISCOVARDovetail_10177 | 13 | 72,843,038 | 72,846,924 | 3886 | hom | no | | | no_coverage | | | | | |
| HG2_PB_SVrefine2Falcon2Bionano_7316 | 14 | 20,550,552 | 20,555,209 | 4657 | hom | no | | | no_coverage | | | | | |
| HG4_PB_SVrefine2Falcon1Dovetail_8950 | 14 | 28,869,843 | 28,872,160 | 2317 | hom | no | | | no_coverage | | | | | |
| HG3_Ill_SVrefine2DISCOVARDovetail_10817 | 14 | 101,335,543 | 101,336,585 | 1042 | hom | no | | | no_coverage | | | | | |
| HG2_Ill_GATKHCSBGrefine_10222 | 15 | 55,218,215 | 55,224,424 | 6209 | hom | yes | DEL | 1 | | 15 | 55218230 | 55224657 | 6427 | both_breakpoints |

| | | | | | | | | | | | | | | |
|---|---|---|---|---|---|---|---|---|---|---|---|---|---|---|
| HG3_Ill_MetaSV_1773 | 15 | 68,426,002 | 68,428,927 | 2925 | hom | yes | DEL | 1 | | 15 | 68426009 | 68429129 | 3120 | both_breakpoints |
| HG4_PB_SVrefine2Falcon1Dovetail_9461 | 15 | 72,385,956 | 72,388,149 | 2193 | hom | no | | | no_coverage | | | | | |
| HG3_PB_SVrefine2Falcon1Dovetail_9416 | 15 | 76,884,596 | 76,896,938 | 12342 | hom | no | | | no_coverage | | | | | |
| HG2_PB_SVrefine2PB10Xhap12_11265 | 15 | 83,551,620 | 83,557,670 | 6050 | hom | no | | | no_coverage | | | | | |
| HG4_PB_SVrefine2Falcon1Dovetail_9558 | 15 | 99,574,407 | 99,575,533 | 1126 | hom | yes | DEL | 1 | | 15 | 99574471 | 99575859 | 1388 | both_breakpoints |
| HG2_Ill_MetaSV_1764 | 16 | 16,934,363 | 16,940,414 | 6051 | hom | yes | DEL | 1 | | 16 | 16934255 | 16940493 | 6238 | both_breakpoints |
| HG2_PB_SVrefine2Falcon2Bionano_7851 | 16 | 18,832,524 | 18,838,376 | 5852 | hom | yes | DEL | 1 | | 16 | 18832397 | 18838387 | 5990 | both_breakpoints |
| HG2_PB_SVrefine2PBcRplusDovetail_4160 | 17 | 16,390 | 17,494 | 1104 | hom | no | | | no_SV_call | | | | | |
| HG2_PB_SVrefine2PBcRplusDovetail_4161 | 17 | 20,542 | 21,552 | 1010 | hom | no | | | no_coverage | | | | | |
| HG2_Ill_SVrefine2DISCOVARplusDovetail_4936 | 17 | 136,688 | 138,370 | 1682 | hom | no | | | no_coverage | | | | | |
| HG2_Ill_SVrefine2DISCOVARplusDovetail_4940 | 17 | 193,768 | 197,057 | 3289 | hom | yes | DEL | 1 | | 17 | 193604 | 197062 | 3458 | both_breakpoints |
| HG4_PB_SVrefine2Falcon1Dovetail_10290 | 17 | 14,189,910 | 14,191,555 | 1645 | hom | yes | DEL | 1 | | 17 | 14189654 | 14191556 | 1902 | both_breakpoints |
| HG3_Ill_SVrefine2DISCOVARDovetail_11844 | 17 | 18,791,138 | 18,792,163 | 1025 | hom | no | | | no_coverage | | | | | |
| HG3_Ill_SVrefine2DISCOVARDovetail_12063 | 17 | 68,455,086 | 68,461,176 | 6090 | hom | yes | DEL | 1 | | 17 | 68454947 | 68461177 | 6230 | both_breakpoints |
| HG2_PB_SVrefine2PBcRplusDovetail_4548 | 17 | 78,461,681 | 78,463,194 | 1513 | hom | no | | | no_coverage | | | | | |
| HG3_PB_SVrefine2PBcRDovetail_8377 | 18 | 21,903,354 | 21,904,585 | 1231 | hom | no | | | no_coverage | | | | | |
| HG2_PB_SVrefine2Falcon2Bionano_8553 | 18 | 38,864,785 | 38,868,343 | 3558 | hom | no | | | no_coverage | | | | | |
| HG3_PB_SVrefine2PBcRDovetail_8451 | 18 | 51,952,963 | 51,956,959 | 3996 | hom | no | | | no_coverage | | | | | |
| HG2_PB_SVrefine2Falcon1plusDovetail_5298 | 18 | 74,104,663 | 74,106,108 | 1445 | hom | no | | | no_coverage | | | | | |
| HG3_Ill_GATKHCSBGrefine_11465 | 18 | 75,266,997 | 75,268,159 | 1162 | hom | no | | | no_coverage | | | | | |
| HG2_PB_SVrefine2Falcon2Bionano_8619 | 18 | 76,129,971 | 76,132,889 | 2918 | hom | no | | | no_coverage | | | | | |
| HG2_PB_SVrefine2PBcRplusDovetail_4929 | 18 | 77,309,921 | 77,312,061 | 2140 | hom | no | | | no_coverage | | | | | |
| HG2_PB_SVrefine2PB10Xhap12_12906 | 19 | 566,138 | 569,346 | 3208 | hom | no | | | no_coverage | | | | | |
| HG3_PB_SVrefine2Falcon1Dovetail_11072 | 19 | 3,172,555 | 3,174,975 | 2420 | hom | no | | | no_coverage | | | | | |
| HG2_PB_SVrefine2PB10Xhap12_12984 | 19 | 3,972,951 | 3,974,022 | 1071 | hom | no | | | no_SV_call | | | | | |
| HG3_PB_SVrefine2Falcon1Dovetail_11117 | 19 | 12,694,655 | 12,698,713 | 4058 | hom | no | | | no_SV_call | | | | | |
| HG4_Ill_SVrefine2DISCOVARDovetail_12464 | 19 | 14,732,331 | 14,734,128 | 1797 | hom | no | | | no_coverage | | | | | |
| HG2_PB_assemblyticsfalcon_12460 | 19 | 30,386,534 | 30,401,481 | 14947 | hom | no | | | no_coverage | | | | | |
| HG3_Ill_SVrefine2DISCOVARDovetail_12767 | 19 | 30,388,790 | 30,393,100 | 4310 | hom | no | | | no_coverage | | | | | |
| HG4_Ill_SVrefine2DISCOVARDovetail_12645 | 19 | 46,941,767 | 46,943,181 | 1414 | hom | no | | | no_SV_call | | | | | |
| HG3_10X_allpass_3428 | 19 | 51,406,755 | 51,408,331 | 1576 | hom | no | | | no_coverage | | | | | |
| HG2_PB_SVrefine2PB10Xhap12_13339 | 19 | 53,689,169 | 53,691,951 | 2782 | hom | yes | DEL | 1 | | 19 | 53689149 | 53692090 | 2941 | both_breakpoints |
| HG4_Ill_SVrefine2DISCOVARDovetail_12782 | 20 | 2,803,147 | 2,806,409 | 3262 | hom | yes | DEL | 1 | | 20 | 2803069 | 2806441 | 3372 | both_breakpoints |
| HG2_10X_SVrefine210Xhap12_12935 | 20 | 21,285,901 | 21,288,579 | 2678 | hom | no | | | low_MapQ | | | | | |
| HG3_Ill_SVrefine2DISCOVARDovetail_13099 | 20 | 31,310,837 | 31,312,741 | 1904 | hom | no | | | no_coverage | | | | | |
| HG4_PB_SVrefine2PBcRDovetail_9353 | 20 | 32,815,928 | 32,819,225 | 3297 | hom | yes | DEL | 1 | | 20 | 32815817 | 32819226 | 3409 | both_breakpoints |
| HG3_PB_SVrefine2Falcon1Dovetail_11580 | 20 | 42,024,594 | 42,025,792 | 1198 | hom | yes | DEL | 1 | | 20 | 42024506 | 42025820 | 1314 | both_breakpoints |
| HG2_PB_SVrefine2PB10Xhap12_13635 | 20 | 42,271,853 | 42,274,577 | 2724 | hom | no | | | no_coverage | | | | | |
| HG2_PB_SVrefine2Falcon1plusDovetail_7049 | 20 | 54,434,610 | 54,440,616 | 6006 | hom | yes | DEL | 1 | | 20 | 54434435 | 54440636 | 6201 | both_breakpoints |
| HG2_PB_SVrefine2PB10Xhap12_14140 | 21 | 47,388,021 | 47,389,724 | 1703 | hom | no | | | no_coverage | | | | | |
| HG4_PB_assemblyticsfalcon_16180 | 21 | 47,388,030 | 47,397,734 | 9704 | hom | no | | | no_coverage | | | | | |
| HG2_10X_SVrefine210Xhap12_13571 | 22 | 24,195,933 | 24,198,505 | 2572 | hom | yes | DEL | 1 | | 22 | 24195676 | 24198515 | 2839 | both_breakpoints |
| HG4_PB_SVrefine2PBcRDovetail_9803 | 22 | 27,168,066 | 27,169,792 | 1726 | hom | no | | | no_coverage | | | | | |
| HG2_PB_SVrefine2PBcRplusDovetail_7046 | 22 | 37,143,113 | 37,147,839 | 4726 | hom | yes | DEL | 1 | | 22 | 37143098 | 37148008 | 4910 | both_breakpoints |

#### Supplementary Table 2

| chrom | start | end | size_estimate_bp | call_class | Notes | Catalogue and evidence |
|---|---|---|---|---|---|---|
| 1 | 58743887 | 58744960 | 1073 | DEL | Hom | PASS Hom catalogue match (913 bp, <1 kb). Excluded from precision. |
| 1 | 92483059 | 92484211 | 1152 | DEL | Het | PASS Het catalogue match (982 bp, <1 kb). Excluded from precision. |
| 1 | 144896268 | 144906810 | 10542 | DEL | Likely TP | Non-PASS match at both breakpoints. Partial coverage loss and high-quality bulk pairs. |
| 1 | 159648645 | 159649809 | 1164 | DEL | Hom | PASS Hom catalogue match (950 bp, <1 kb). Excluded from precision. |
| 1 | 238852278 | 238853419 | 1141 | DEL | Het | PASS Het catalogue match (943 bp, <1 kb). Excluded from precision. |
| 1 | 243782623 | 243783836 | 1213 | DEL | Hom | PASS Hom catalogue match (997 bp, <1 kb). Excluded from precision. |
| 2 | 89854009 | 89856669 | 2660 | DEL | Likely TP | Non-PASS match at both breakpoints. Partial coverage loss and high-quality bulk pairs. |
| 3 | 189224306 | 189225372 | 1066 | DEL | Het | PASS Het catalogue match (864 bp, <1 kb). Excluded from precision. |
| 3 | 194543212 | 194546302 | 3090 | DEL | FP | Non-PASS catalogue match. Bulk pairs observed. Conservatively counted as FP. |
| 4 | 59428819 | 59435539 | 6720 | DEL | FP | Non-PASS catalogue match. Bulk pairs observed. Conservatively counted as FP. |
| 4 | 147730059 | 147731173 | 1114 | DEL | Het | PASS Het catalogue match (913 bp, <1 kb). Excluded from precision. |
| 4 | 152340671 | 152341674 | 1003 | DEL | Het | PASS Het catalogue match (847 bp, <1 kb). Excluded from precision. |
| 5 | 8937631 | 8938821 | 1190 | DEL | Het | PASS Het catalogue match (973 bp, <1 kb). Excluded from precision. |
| 5 | 125496630 | 125498799 | 2169 | DEL | FP | Non-PASS catalogue match. Bulk pairs observed. Conservatively counted as FP. |
| 5 | 127172266 | 127173307 | 1041 | DEL | Hom | PASS Hom catalogue match (950 bp, <1 kb). Excluded from precision. |
| 5 | 144871830 | 144872954 | 1124 | DEL | Het | PASS Het catalogue match (901 bp, <1 kb). Excluded from precision. |
| 6 | 30993954 | 30995100 | 1146 | DEL | Likely TP | Non-PASS match at both breakpoints. Partial coverage loss and high-quality bulk pairs. |
| 6 | 31356131 | 31453167 | 97036 | DEL | FP | No catalogue match. Bulk pairs observed. Conservatively counted as FP. |
| 6 | 32461254 | 32468821 | 7567 | DEL | Hom | No catalogue match. Coverage supports Hom. Excluded from precision. |
| 6 | 32594333 | 32596473 | 2140 | DEL | FP | Non-PASS catalogue match. Bulk pairs observed. Conservatively counted as FP. |
| 6 | 52651094 | 52652694 | 1600 | DEL | FP | No catalogue match. Bulk pairs observed. Conservatively counted as FP. |
| 6 | 68101463 | 68112240 | 10777 | DEL | FP | Non-PASS catalogue match. Bulk pairs observed. Conservatively counted as FP. |
| 7 | 62002245 | 62004962 | 2717 | DEL | Hom | Non-PASS catalogue match. Coverage supports Hom. Excluded from precision. |
| 7 | 141760416 | 141786988 | 26572 | DEL | FP | Non-PASS catalogue match. Bulk pairs observed. Conservatively counted as FP. |
| 8 | 18651268 | 18652426 | 1158 | DEL | Hom | PASS Hom catalogue match (974 bp, <1 kb). Excluded from precision. |
| 8 | 37050732 | 37051854 | 1122 | DEL | Hom | PASS Hom catalogue match (974 bp, <1 kb). Excluded from precision. |
| 8 | 138742827 | 138743871 | 1044 | DEL | Hom | PASS Hom catalogue match (896 bp, <1 kb). Excluded from precision. |
| 9 | 107596077 | 107597172 | 1095 | DEL | Het | PASS Het catalogue match (916 bp, <1 kb). Excluded from precision. |
| 10 | 95545424 | 95546608 | 1184 | DEL | Hom | Non-PASS catalogue match. Coverage supports Hom. Excluded from precision. |
| 10 | 97206949 | 97208027 | 1078 | DEL | FP | Non-PASS catalogue match. Bulk pairs observed. Conservatively counted as FP. |
| 10 | 127191560 | 127197236 | 5676 | DEL | FP | Non-PASS catalogue match. Bulk pairs observed. Conservatively counted as FP. |
| 11 | 29967533 | 29968551 | 1018 | DEL | Het | PASS Het catalogue match (888 bp, <1 kb). Excluded from precision. |
| 11 | 48600658 | 48604291 | 3633 | DEL | Hom | Non-PASS catalogue match. Coverage supports Hom. Excluded from precision. |
| 11 | 50640076 | 50641133 | 1057 | DEL | Hom | Non-PASS catalogue match. Coverage supports Hom. Excluded from precision. |
| 11 | 54885193 | 54892327 | 7134 | DEL | FP | No catalogue match. Bulk pairs observed. Conservatively counted as FP. |
| 11 | 129013515 | 129015318 | 1803 | DEL | FP | Non-PASS catalogue match. Bulk pairs observed. Conservatively counted as FP. |
| 12 | 5082178 | 5083178 | 1000 | DEL | FP | Non-PASS catalogue match. Bulk pairs observed. Conservatively counted as FP. |
| 12 | 7951098 | 8058285 | 107187 | DEL | FP | No catalogue match. Bulk pairs observed. Conservatively counted as FP. |
| 12 | 63939035 | 64134212 | 195177 | DEL | FP | No catalogue match. Bulk pairs observed. Conservatively counted as FP. |
| 13 | 29786932 | 29922251 | 135319 | DEL | FP | No catalogue match. Bulk pairs observed. Conservatively counted as FP. |
| 13 | 56851593 | 56852680 | 1087 | DEL | Het | PASS Het catalogue match (830 bp, <1 kb). Excluded from precision. |
| 13 | 74915200 | 74916776 | 1576 | DEL | FP | Non-PASS catalogue match. Bulk pairs observed. Conservatively counted as FP. |
| 14 | 106932595 | 107174957 | 242362 | DEL | FP | Non-PASS catalogue match. Bulk pairs observed. Conservatively counted as FP. |
| 16 | 75429637 | 75430737 | 1100 | DEL | Hom | PASS Hom catalogue match (842 bp, <1 kb). Excluded from precision. |
| 17 | 17230220 | 17231308 | 1088 | DEL | Het | PASS Het catalogue match (876 bp, <1 kb). Excluded from precision. |
| 17 | 46615777 | 46617329 | 1552 | DEL | FP | Non-PASS catalogue match. Bulk pairs observed. Conservatively counted as FP. |
| 17 | 78575276 | 78576379 | 1103 | DEL | Hom | PASS Hom catalogue match (884 bp, <1 kb). Excluded from precision. |
| 19 | 21902874 | 21905562 | 2688 | DEL | Hom | Non-PASS catalogue match. Coverage supports Hom. Excluded from precision. |
| 19 | 27801342 | 27806318 | 4976 | DEL | FP | No catalogue match. Bulk pairs observed. Conservatively counted as FP. |
| 19 | 46278518 | 46279617 | 1099 | DEL | Hom | Non-PASS catalogue match. Coverage supports Hom. Excluded from precision. |
| 20 | 26297487 | 26302772 | 5285 | DEL | FP | Non-PASS catalogue match. Bulk pairs observed. Conservatively counted as FP. |
| 20 | 61724662 | 61725674 | 1012 | DEL | Het | PASS Het catalogue match (898 bp, <1 kb). Excluded from precision. |
| 22 | 24273991 | 24311298 | 37307 | DEL | FP | Non-PASS catalogue match. Bulk pairs observed. Conservatively counted as FP. |
| 22 | 31419740 | 31420854 | 1114 | DEL | Het | PASS Het catalogue match (935 bp, <1 kb). Excluded from precision. |
| | | | | | | Notes key |
| | | | | | | Hom (16): 9 PASS Hom matches <1 kb and 7 additional coverage-supported Hom calls. |
| | | | | | | Het (13): PASS heterozygous catalogue matches <1 kb. |
| | | | | | | Likely TP (3): likely true deletions with unresolved genotype. |
| | | | | | | All 32 Hom, Het and Likely TP calls are excluded from the TP and FP counts for precision. |
| | | | | | | FP (22): conservatively scored false positives; 15 match non-PASS catalogue entries. |
| | | | | | | Full catalogue: 46 of 54 calls matched, comprising 22 PASS and 24 non-PASS matches. |
| | | | | | | Precision: 96 / (96 + 22) = 81.4%. |
